# ParB C-terminal lysine residues are essential for dimerization, *in vitro* DNA sliding and *in vivo* function

**DOI:** 10.1101/2025.06.10.659001

**Authors:** Aleksey Aleshintsev, Linsey E. Way, Bianca Guerra, Lois Akosua Serwaa, Miranda Molina, Xindan Wang, HyeongJun Kim

## Abstract

The broadly conserved ParB protein performs crucial functions in bacterial chromosome segregation and replication regulation. The cellular function of ParB requires it to dimerize, recognize *parS* DNA sequences, clamp on DNA, then slide to adjacent sequences through nonspecific DNA binding. How ParB coordinates nonspecific DNA binding and sliding remains elusive. Here, we combine multiple *in vitro* biophysical and computational tools and *in vivo* approaches to address this question. We found that the five conserved lysine residues in the C-terminal domain of ParB play distinct roles in proper positioning and sliding on DNA, and their integrity is crucial for ParB’s *in vivo* functions. Many proteins with diverse cellular activities need to move on DNA while loosely bound. Our findings reveal the detailed molecular mechanism by which multiple flexible basic residues enable DNA binding proteins to efficiently slide along DNA.

## Introduction

Accurate chromosome replication and segregation are essential for all living organisms. For most bacterial chromosomes, this task is performed by a well-conserved ParABS system. ParA is a Walker-type ATPase that binds non-specifically to DNA (1, 2); ParB is a CTPase that binds to the centromere-like *parS* DNA sequences, which are present as one or multiple copies in the vicinity of the origin of replication (3–5). ParB stimulates the ATP hydrolysis of ParA, and repeated cycles of ParA and ParB interactions promote the segregation of the newly replicated origins (6–9). ParB plays a separate function in loading the structural maintenance of chromosomes (SMC) complexes to the chromosome, which is also important for chromosome segregation (10, 11). Finally, ParA’s ATPase cycle regulates the initiation of DNA replication (12). Therefore, the ParABS system plays an instrumental role in the segregation and replication of bacterial chromosomes (13, 14).

In some bacterial species, such as *Caulobacter crescentus*, ParB (and ParA) proteins are essential for cell viability (15). In *Bacillus subtilis*, cells lacking *parB (*also known as *spo0J*) are viable in vegetative growth but exhibit a 50-100 fold increase in anucleate cells (16), emphasizing its function in the faithful chromosome segregation. In addition, ParB is required for sporulation, which is a developmental procedure *B. subtilis* undergoes under starvation and other stresses.

ParB proteins have three conserved structural domains (Extended Data Fig. 1a). The N-terminal domain (NTD) (or the ParB/Srx domain) contains cytidine triphosphate (CTP) binding motifs (17–20) and ParA interaction domains (1, 21, 22). A recent study implies that NTD also mediates ParB multimerization and the formation of phase-separated condensate (23). The central DNA-binding domain (CDBD) specifically recognizes *parS* DNA through its helix-turn-helix motif (19, 20, 24–27). The C-terminal domain (CTD) is essential for nonspecific DNA binding (28, 29), ParB dimerization (20, 24, 30), and DNA condensation via ParB-ParB bridging interactions (29). Notably, a CTD nuclear magnetic resonance (NMR) structure (PDB: 5NOC) revealed a surface-exposed lysine-rich DNA-binding patch (29).

ParB proteins are found not only from their cognate *parS* DNA binding sites but also in the 10-20 kb vicinity of each *parS* site (4, 30–32), a phenomenon termed spreading. Earlier models proposed a 1D filament-like spreading (4, 30, 32) where ParB polymerizes along DNA to form a continuous filament after loading on *parS*. This model was backed up by the observation of ParB spreading being attenuated by a protein “roadblock” positioned on one side of *parS* (32). However, quantitative immunoblots and fluorescence microscopy in *B. subtilis* estimated that the 1D filament model alone can account for only ∼500 bp (instead of 10-20 kb) of spreading due to the low ParB concentration (∼20 per *parS* site) (33). The authors demonstrated that ParB proteins are capable of compacting flow-stretched DNA and proposed that DNA bridging events are required for ParB spreading (33). Later, Sanchez and coworkers discovered that a majority of ParB molecules are localized around *parS* and proposed a nucleation and caging model (34). The paradigm-shifting discoveries that ParB proteins bind to and hydrolyze cytidine triphosphate (CTP) (18, 35, 36) established models involving ParB sliding on DNA (18, 20). Here, a ParB dimer adopts an open NTD conformation (the NTD-gate) and binds to the *parS* DNA sequence through the helix-turn-helix motif on the CDBD (the DNA-gate). CTP binding self-dimerizes the NTD-gate, closing the DNA-gate and forming a DNA sliding clamp (19, 20, 35, 36). The NTD- and DNA-gates closures release *parS* DNA into a compartment between the DNA-gate and CTD, allowing ParB to slide away from *parS* (37, 38). CTP hydrolysis reopens the gates, causing ParB to be unloaded and recycled (19, 20, 35).

In *B. subtilis*, the CTD contains five lysines (K252, K255, K256, K257, and K259). Mutating a subset of (K255A-K257A or K252A-K255A-K259A) diminished DNA condensation in a magnetic tweezer-based assay (29). The results indicate that the positively charged lysine residues directly interact with the negatively charged DNA backbone. Thus, it is puzzling how the ParB clamp slides while its CTD interacts with DNA. The understanding of ParB CTD dynamics and its effects on CTD functions is limited. In this study, we employed NMR, single-molecule, biochemical approaches, molecular dynamics simulation, cellular fluorescence imaging, and sporulation initiation assay to investigate the molecular mechanism and cellular function of CTD dynamics. We demonstrate that each CTD lysine has a distinct role. Fast lysine dynamics, occurring on timescales of picoseconds to nanoseconds, promote efficient DNA sliding. We also show that any single lysine mutation substantially impacts DNA bridging *in vitro* and argue that the integrity of these five lysines is crucial for positioning DNA properly in the ParB sliding clamp. Our data show that CTD dimerization is important for ParB foci formation and sporulation initiation. Additionally, we provide insights into how DNA-binding proteins in diverse protein families use multiple basic residues for fast dynamics.

## Results

### NMR parameters (^15^N R1, R2, and NOE) of apo CTD indicate fast backbone dynamics in the lysine residues

The mobility of the protein backbone chain is important for its function. For the *B. subtilis* ParB (BsParB) protein, the CTD is a nonspecific DNA-binding domain (29), and we hypothesize that its backbone dynamics promote DNA sliding. To understand the CTD backbone dynamics upon nonspecific DNA binding, we performed nuclear magnetic resonance (NMR) relaxation studies for ParB CTD(217–282) expressed in *Escherichia coli* (Extended Data Fig. 1b). As a starting point, building upon previously reported residue assignments of apo-CTD (29), we analyzed 57 non-overlapping peaks. These assignments are presented on a ^1^H-^15^N-heteronuclear single quantum coherence (HSQC) spectrum of CTD at 308 K, pH 6.1 (Extended Data Fig. 2a).

Following the residue assignments, to study the protein motion of apo CTD for each backbone amide on a fast (picoseconds to nanoseconds) time scale, we characterized three relaxation parameters: the self-relaxation rates (^15^N R1 and R2) and the steady-state heteronuclear Overhauser effect ({^1^H}^15^N–NOE) that corresponds to dipolar cross-relaxation between two coupled ^1^H and ^15^N spins. R1 and R2 are rate constants with a unit of s^-1^ that characterize the decay rates of two types of magnetizations: longitudinal and transverse. These relaxation rates depend on how observed nuclei, such as ^1^H and ^15^N, experience specific oscillating magnetic fields. In proteins, these specific oscillating fields are due to the movements of magnetic nuclei in the overall bulk magnetic field of the NMR magnet or due to relative movements with respect to each magnetic field (39). Thus, relaxation results reflect protein motion and provide insightful information. Apo CTD’s relaxation data (R1, R2, and NOE) were acquired at an 11.7 T field (500 MHz). All relaxation parameters were plotted as a function of the amino acid residues (Fig. 1a-d and Table 1.)

**Fig. 1.**
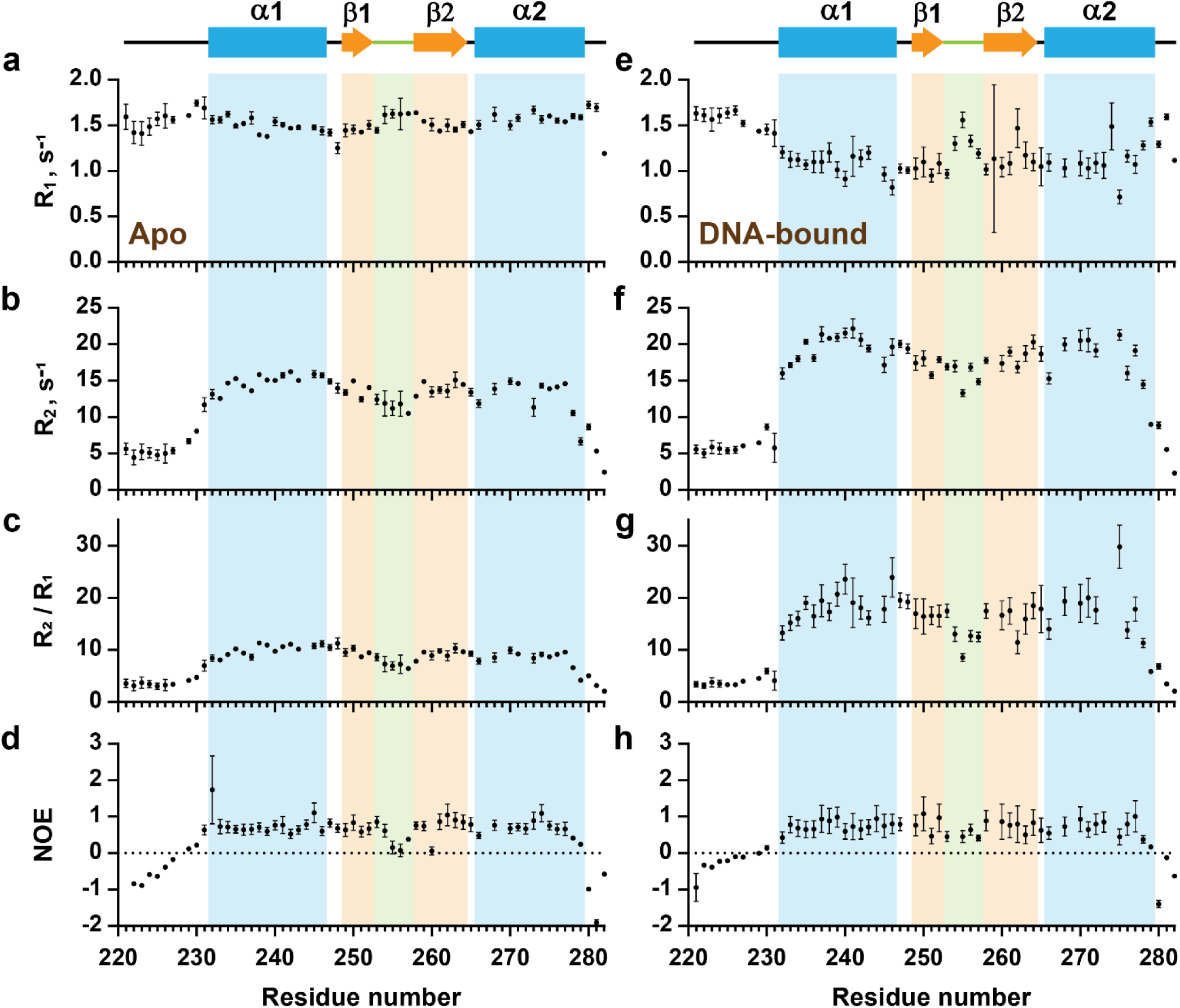
Relaxation data of apo and DNA-bound CTD at 11.7 T. 400 µM CTD in PBS buffer, pH 6.1 at 308 K. For apo CTD: (**a**) R1. (**b**) R2. (**c**) R2/R1. (**d**) NOE. For DNA-bound CTD: (**e**) R1. (**f**) R2. (**g**) R2/R1. (**h**) NOE. The secondary structure of the apo CTD determined from the NMR structure (PDB entree 5NOC) is displayed at the top: α-helices (cyan bars), β-strands (light orange arrows), loops (black lines), and linker region (light green).

**Table 1:** Average relaxation parameters per secondary structures and terminal loops (top half) and average order parameters, S^2^ per secondary structures, and terminal loops in DNA-bound CTD (bottom half) in apo and DNA-bound CTD.

| Apo CTD |  |  |  |  |  |  |  |  |  |
| --- | --- | --- | --- | --- | --- | --- | --- | --- | --- |
| | N-Term<br>(221-230) | α-helix 1<br>(231-246) | Loop 1<br>(247-248) | β-strand 1<br>(249-252) | Linker<br>(253-257) | β-strand 2<br>(258-264) | Loop 2<br>(265) | α-helix 2<br>(266-279) | C-term<br>(280-282) |
| R<sub>1</sub> | 1.556 | 1.516 | 1.337 | 1.459 | 1.589 | 1.514 | 1.433 | 1.576 | 1.538 |
| R<sub>2</sub> | 5.610 | 14.664 | 14.441 | 13.725 | 11.578 | 14.027 | 13.414 | 12.805 | 5.499 |
| R<sub>2</sub>/R<sub>1</sub> | 3.612 | 9.781 | 10.870 | 9.501 | 7.307 | 9.278 | 9.301 | 8.301 | 3.518 |
| NOE | -0.396 | 0.702 | 0.755 | 0.682 | 0.416 | 0.746 | 0.786 | 0.669 | -1.159 |
| DNA-bound CTD |  |  |  |  |  |  |  |  |  |
| R<sub>1</sub> | 1.570 | 1.103 | 1.018 | 1.040 | 1.270 | 1.146 | 1.047 | 1.137 | 1.335 |
| R<sub>2</sub> | 6.038 | 18.607 | 19.731 | 17.291 | 15.789 | 18.332 | 18.688 | 15.928 | 5.584 |
| R<sub>2</sub>/R<sub>1</sub> | 3.880 | 17.355 | 19.388 | 16.628 | 12.845 | 16.261 | 17.853 | 16.855 | 4.142 |
| NOE | -0.240 | 0.746 | 0.777 | 0.818 | 0.587 | 0.724 | 0.624 | 0.664 | -0.717 |
| Apo CTD |  |  |  |  |  |  |  |  |  |
| | N-Term<br>(229-230) | α-helix 1<br>(231-246) | Loop 1<br>(247-248) | β-strand 1<br>(249-252) | Linker<br>(253-257) | β-strand 2<br>(258-264) | Loop 2<br>(265) | α-helix 2<br>(266-279) | C-term<br>(280-282) |
| S<sup>2</sup> | 0.46 | 0.91 | 0.87 | 0.89 | 0.77 | 0.90 | 0.87 | 0.84 | 0.13 |
| S<sup>2</sup> err | 0.03 | 0.02 | 0.02 | 0.02 | 0.04 | 0.02 | 0.01 | 0.02 | 0.03 |
| DNA-bound CTD |  |  |  |  |  |  |  |  |  |
| S<sup>2</sup> | 0.31 | 0.85 | 0.89 | 0.84 | 0.77 | 0.88 | 0.90 | 0.75 | 0.21 |
| S<sup>2</sup> err | 0.02 | 0.03 | 0.02 | 0.03 | 0.02 | 0.03 | 0.05 | 0.04 | 0.01 |
| S<sup>2</sup> | 0.31 | 0.85 | 0.89 | 0.84 | 0.77 | 0.88 | 0.90 | 0.75 | 0.21 |

The R2/R1 ratio can estimate the overall molecular tumbling correlation time, τm. This estimation is achieved when conformational exchanging motions are absent and internal motions are fast, allowing the R2/R1 ratio to depend only on the overall rotational correlation time (τm) (40, 41). Therefore, for our τm calculations, we excluded residues with low NOE that correspond to a large amplitude of internal motions and are indicative of a timescale of motions longer than hundreds of picoseconds (42). Residues having a greater than 1.0 SD (standard deviation) for the R2/R1 ratio were also excluded from the calculations due to the possible effect of the additional contribution of conformational exchange Rex to R2 (40, 43). The initial mean correlation time (τm) was 11.02±0.66 (S.D.) ns, and the mean R2/R1 ratio equals 7.92±2.65 (S.D.). (See Methods). R2/R1 for the apo CTD shows different dynamics between secondary structures along the backbone. Secondary structures on the N-terminal half of the CTD have higher than average R2/R1 (9.079), suggesting intermediate timescale motion. On average, the N-terminal half exhibits larger R2/R1 values compared to the secondary structures of the C-terminal half of the CTD (Fig. 1c). The low average R2/R1 values of the C-terminal α-helix and especially the linker with its DNA binding lysine residues suggest mostly fast timescale backbone dynamics of these regions in the apo CTD (Fig. 1c). The apo relaxation data indicate distinct dynamics between different secondary structures and reveal highly dynamic linker regions.

### Determining apo CTD’s dynamic parameters (S^2^, Rex, τ) with Model-Free analysis

R2/R1 data and data from pdbinertia (See Methods) were further used to estimate rotational diffusion tensors (for overall tumbling motion) and to determine the appropriate motional model (spherical, axially symmetric, or fully anisotropic). The results of the quadric_diffusion program (See Methods) were collected in Supplementary Table 1. The model of molecular tumbling was selected with the smallest chi-square (χ^2^) and an F value. For the apo CTD, the overall tumbling is fully anisotropic with anisotropy DA = 2Dzz/(Dxx+Dyy) equal to 1.282±0.043 based on comparing χ^2^ and F statistics for each model. The average overall rotational correlation time, τm, equals (6×Diso)^-1^ = 11.08±0.06 ns. We should note that the degree of anisotropy is small, so we can treat the tumbling of apo CTD as axially symmetric with the FAST-Modelfree program applying output pdb file for anisotropic diffusion analysis. We applied initial estimates of τm, Dpar/Dper, θ and φ, and other relaxation parameters in the FAST-Modelfree program (See Methods) using the axially symmetric model for rotational diffusion tensor. The program then iteratively refined parameters such as S^2^ (order parameter) and τe to fit ^15^N relaxation data. Five spectral density functional models fit the data (Supplementary Table 2). The results for rotational diffusion tensor show that the relaxation of NH vectors for the majority of the apo CTD residues fit the simplest model, Model 1 (29 residues), in which S^2^ is the only parameter to fit. Models 2, 3, 4, and 5 fit three, nine, three, and seven residues, respectively. No model was assigned to E281. The resulting parameters are represented in the Supplementary Table 3. Twelve residues exhibit additional conformational exchange represented by the Rex term ranging from 0.787±0.258 s^-1^ (L277) to 4.598±0.721 s^-1^ (R280) (Fig. 2a and Supplementary Table 3). Rex can indicate slow conformational changes in the protein. Compared to R2 data, the Rex term of most residues may be considered low except for R280, located in the C-terminal loop. Nine residues with Rex are located in the N-terminal and C-terminal α-helices, suggesting these regions undergo specific conformational changes on a slow time scale, especially on α-helix 1. The results of Model-Free analysis suggest that CTD backbone does not experience extended conformational changes in the apo state while having mostly axially symmetric molecular tumbling.

**Fig. 2.**
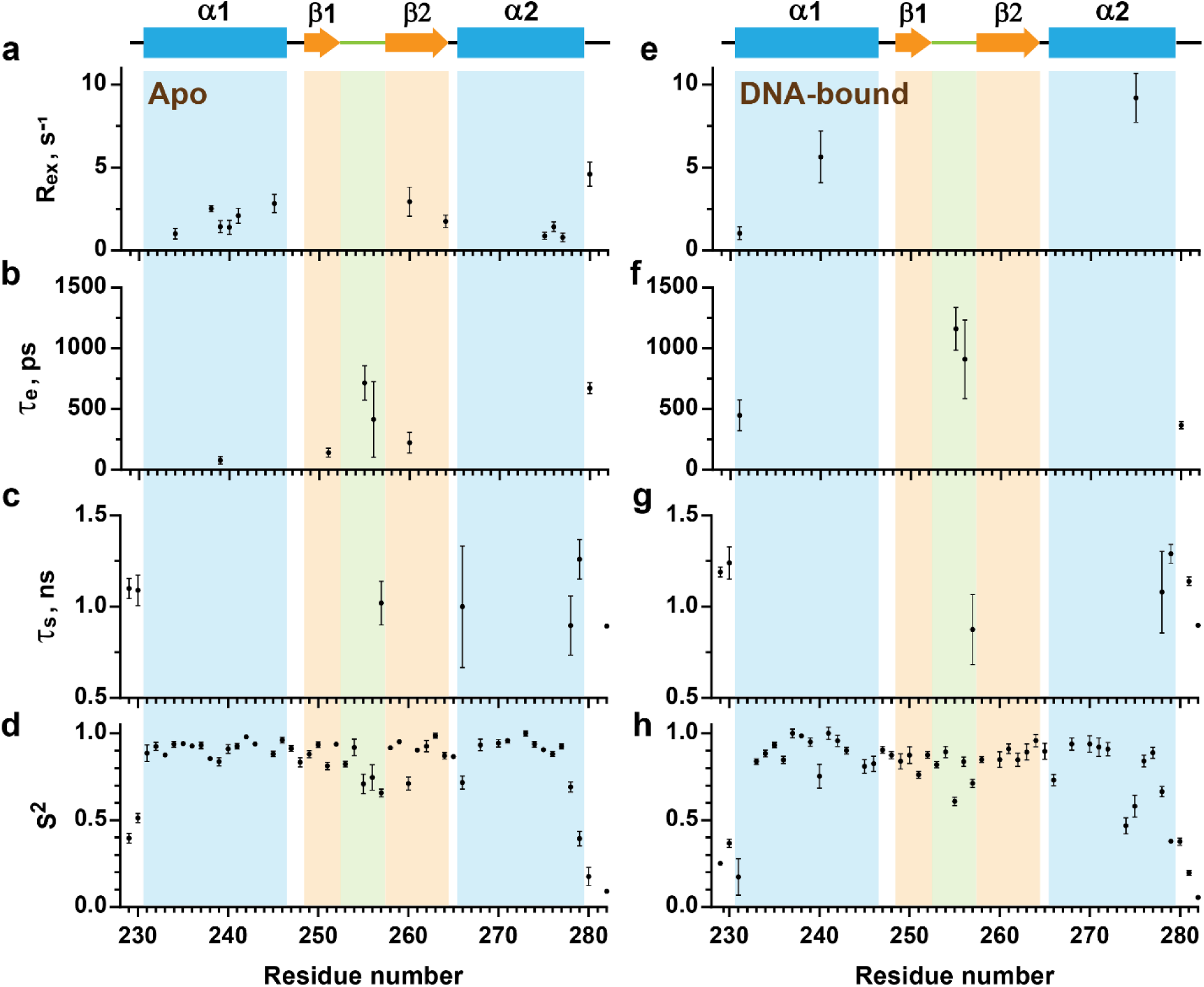
Parameters defining the backbone dynamics of apo and DNA-bound CTD as a function of the residue number. **a-d**, Backbone dynamics parameters for apo CTD. Rex, τe, τs, and S^2^ are the chemical exchange contribution, the internal correlation time, residue with internal correlation times at slower timescales, and the order parameter, respectively. The secondary structure determined from the NMR structure (PDB entree 5NOC) is displayed at the top: α-helices (cyan bars), β-strands (light orange arrows), loop regions (black lines), and linker region (light green). **e-h**, Backbone dynamics parameters for DNA-bound CTD.

### S^2^, τs, τe distribution of apo CTD reveals contrasting motions in the DNA-binding lysine residues

Most residues of apo CTD exhibit fast internal motion (τe ≪ τm), and only the order parameter S^2^ was needed to fit Model 1 (Supplementary Table 3). Only thirteen residues’ NH vectors needed the inclusion of internal correlation time parameters, τe and τs for their relaxation model calculations (Supplementary Table 3 and Fig. 2b,c). Figure 2b shows that the fast timescale (τe) motions were detected in the linker region (K255 and K256) and the surrounding β-strands. Relatively slower internal motions (τs) were detected predominantly in the N- and C-terminal residues but also in the linker region (K257) (Fig. 2c).

The order parameter values for angular motion (S^2^) extracted from the Lipari-Szabo’s model-free formalism (44) carry substantial weight in understanding ^1^H-^15^N bond vector internal motions (45). Figures 2d, 3a, and Extended Data Figures 3a,c display the S^2^ values where 0 and 1 represent complete flexibility and rigidity, respectively. The overall average value of S^2^ for apo CTD is 0.83±0.02. The average values of S^2^ for the secondary structural elements and termini regions in the apo CTD are listed in Table 1.

26 NH vectors in apo CTD exhibit S^2^ ≥ 0.9, of which six vectors have S^2^ ≥ 0.95. A high degree of motion restriction was observed in α-helix 1 and both β-strands. In α-helix 2 (266–279), its last two residues, S278 and E279, show highly elevated mobility (low S^2^) for their NH vectors (Fig. 2d), in contrast to the significantly restricted dynamics of the rest of α-helix 2. Both relaxation data (R2/R1 and NOE) and S^2^ values for S278 and E279 show similar trends for the C-terminal end (280–282) (Fig. 1c,d), suggesting that S278 and E279 might not belong to α-helix 2. Apart from these two residues, the general trend for the apo CTD is that both helices are the least dynamic (highly restricted motions), followed by the β-strands.

In contrast, outside of termini regions where the lowest average S^2^ values are expected, the average value of S^2^ is lowest (most flexible) for the linker loop (253–257) (0.77±0.04), where putative DNA-binding lysine residues are positioned (Fig. 2d, 3a, and Extended Data Fig. 3a,c), K257 exhibits a particularly low S^2^ value (0.657). In summary, lysine residues (K255-K257) on the flexible linker region show high flexibility (low S^2^) while those on rigid β-strands (K252 and K259 on β-strands 1 and 2, respectively) show high rigidity (high S^2^). It suggests distinct, location-dependent roles for each lysine residue, as we discuss later.

**Fig. 3.**
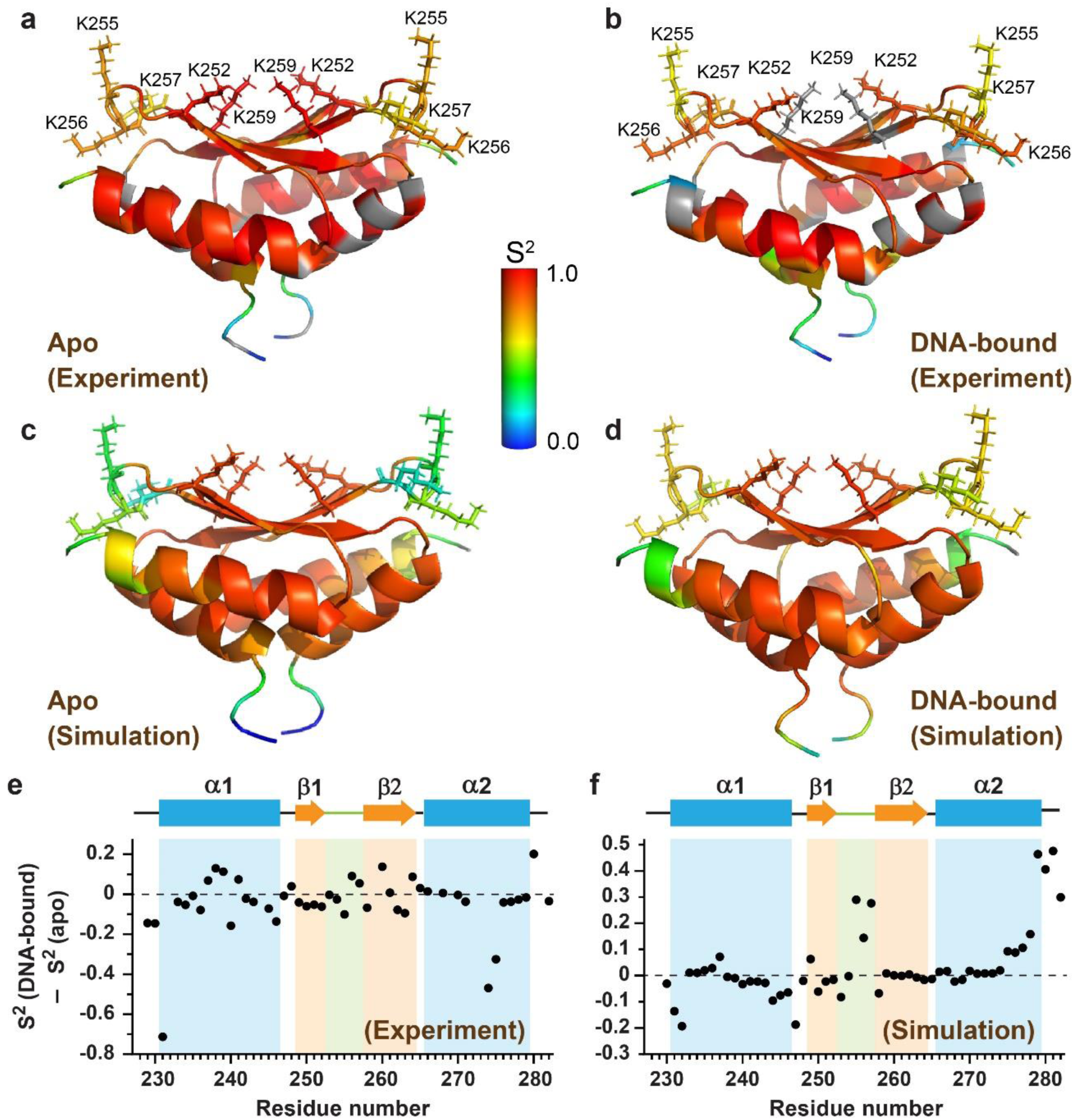
Calculated S^2^ values and their differences upon DNA binding. **a-b**, Calculated S^2^ values from NMR relaxation experiments for (**a**) apo and (**b**) DNA-bound CTD, respectively. The color-coded values are mapped onto CTD structure (PDB entry: 5NOC). Lysines of interest are labeled. Color scheme: gradient from blue to red for residues with S^2^ between 0.0 and 1.0, and gray for residues with no assigned S^2^. **c-d**, Those from molecular dynamics (MD) simulations for (**c**) apo and (**d**) DNA-bound CTD, respectively. **e-f**, Calculated S^2^ value differences from (**e**) NMR experiment data and (**f**) MD simulation data, respectively. Positive and negative values imply enhanced and decreased rigidity, respectively, upon DNA binding.

### Relaxation data show altered backbone dynamics upon DNA binding

Since positively charged amino acid patches on CTD have been known to bind DNA nonspecifically (29), we investigated structural aspects of DNA-bound CTD using NMR relaxation data acquired in the same manner as for the apo CTD. To determine the shift of signals, we titrated CTD with DNA in 0.25 equivalent steps and collected ^1^H-^15^N HSQC for each step (Extended Data Fig. 2b). Several peaks were excluded from the relaxation data due to significant line broadening or peak overlap. All relaxation parameters were plotted as a function of the amino acid residues and are reported in Fig. 1e-h. (See also Table 1.) The initial mean correlation time (τm) was 15.45±1.10 (S.D.) ns, and the mean R2/R1 ratio was 13.84±6.60 (S.D.). Calculations were performed as described in the Methods section. While R1 of the apo CTD exhibits similar values across the chain, the R1 data of the DNA-bound CTD shows a marked decrease for the secondary structures (Fig. 1a,e). On the other hand, the R2 of the DNA-bound CTD exhibits a significant increase in its values along the secondary structures (Fig. 1b,f). R2/R1 for the DNA-bound CTD significantly increases along the whole chain except for the termini (Fig. 1g). This indicates a slowing down of molecular tumbling due to DNA binding. While we observed the global increase in R2/R1 by adding DNA to the CTD, the R2/R1 patterns in the DNA-bound CTD are somewhat different compared to the apo CTD. Upon binding to the DNA, both β-strands, which participate in the DNA-binding through K252 and K259, have R2/R1 values that are lower or almost the same as the average R2/R1 (16.600), while in the apo-CTD, the R2/R1 of the β-strands are above the average, indicating dynamic changes introduced by DNA. Additionally, both helices in the DNA-bound CTD exhibit higher than average R2/R1 ratios, indicating slower dynamics. However, in the apo state, the C-terminal α-helix’s R2/R1 is below average. Only the dynamics of both loops and the linker are the same as in the apo CTD; the loops, especially the first, have slow motion, while the linker has fast dynamics. These data indicate that DNA introduces dynamic changes to the CTD, and the linker enables DNA recognition.

### Determining DNA-bound CTD’s dynamic parameters (S^2^, Rex, τ) with Model-Free analysis

Despite no DNA-bound structure of CTD being presently available, we decided to evaluate the dynamics of the DNA-bound CTD using the available apo CTD structure (PDB: 5NOC). As in the case of the apo CTD relaxation data analysis above, we determined a motional model of DNA-bound CTD (Supplementary Table 4). The molecular tumbling of DNA-bound CTD was best described as fully anisotropic with DA equal to 1.319±0.074 (χ^2^=22.83, F=1.69). The average overall rotational correlation time, τm, was 15.23±0.23 ns. Compared to apo CTD, the degree of anisotropy of DNA-bound CTD is more considerable. Nonetheless, this anisotropy is still not large, so we can treat the tumbling of DNA-bound CTD as axially symmetric, applying the output pdb file for anisotropic diffusion analysis. The results of using axially symmetric models for rotational diffusion tensors show that the relaxation of NH vectors of the majority of the DNA-bound CTD residues fit the simplest Model 1 (36 residues) (Supplementary Table 5). Two residues were not assigned (A232 and I273). In contrast to apo CTD, only three residues (D231, Y240, and E275) exhibit additional conformational exchange represented by Rex term ranging from 1.037±0.383 s^-1^ (D231) to 9.202±1.468 s^-1^ (E275) (Fig. 2e and Supplementary Table 5). The results of Model-Free analysis suggest that the addition of DNA decreases conformational changes of CTD backbone in comparison to apo CTD.

### S^2^, τs, τe distribution of DNA-bound CTD reveals changes in the dynamics of DNA-binding lysine residues upon addition of DNA

As with apo CTD, the majority of residues in the DNA-bound CTD exhibit fast internal motion and require only S^2^ to fit Model 1 (Supplementary Table 5). The eleven residues’ NH vectors require the inclusion of additional internal correlation time parameters, τe and τs, for their relaxation model calculations (Supplementary Table 5). Comparisons between Fig. 2b,f show that the fast timescale (τe) internal motion becomes slower in the linker region (K255 and K256) when DNA is bound. As in apo CTD, relatively slower internal motions (τs) were detected predominantly in the N- and C-terminal residues but also in the linker region (K257) (Fig. 2g).

Figures 2h, 3b, and Extended Data Figures 3b,d display the order parameter for angular motion, S^2^. The overall average value of S^2^ is 0.76±0.03. (See also Table 1.) Outside of termini regions, the average values of S^2^ are smallest for α-helix 2 (0.75±0.04) and the linker (253–257) (0.77±0.02) (Fig. 2h, 3b, and Extended Data Fig. 3b,d). DNA binding resulted in S^2^ value changes throughout the CTD structure (Fig. 3e). Introducing DNA has prominent consequences on S^2^ values, including (1) more restricted internal motion (higher S^2^) in the middle of α-helix 1 and decreased S^2^ on both of its ends, (2) increased flexibility (lower S^2^) on β-helix 1, and (3) lower and higher rigidities (lower and higher S^2^) for K255 and K256-K257 linker lysine residues, respectively. These results suggest different roles of K255 and K256/K257 in DNA recognition, which require further investigation.

### E261-mediated salt bridges hold two CTD monomers together

The flexible linker region contains three putative DNA-binding lysine residues (K255-K257). We claim that the elevated dynamics of the linker are essential for DNA recognition. Other putative DNA-binding lysine residues (K252 and K259) are located on highly rigid (high S^2^) β-strands. We questioned whether there were any structural grounds for the high rigidity of K252 and K259 residues and investigated surrounding residues from the known structure of apo *B. subtilis* ParB CTD (PDB: 5NOC) (29). We discovered that negatively-charged E261 on β-strand 2 forms salt bridges with K252 on the same CTD monomer and K259 on the other monomer (Fig. 4a). These salt bridges confer the high rigidity of the K252 and K259 residues and imply different roles for lysines within β-strands (K252 and K259) compared to those within the linker region (K255-K257) (See Discussion.)

**Fig. 4.**
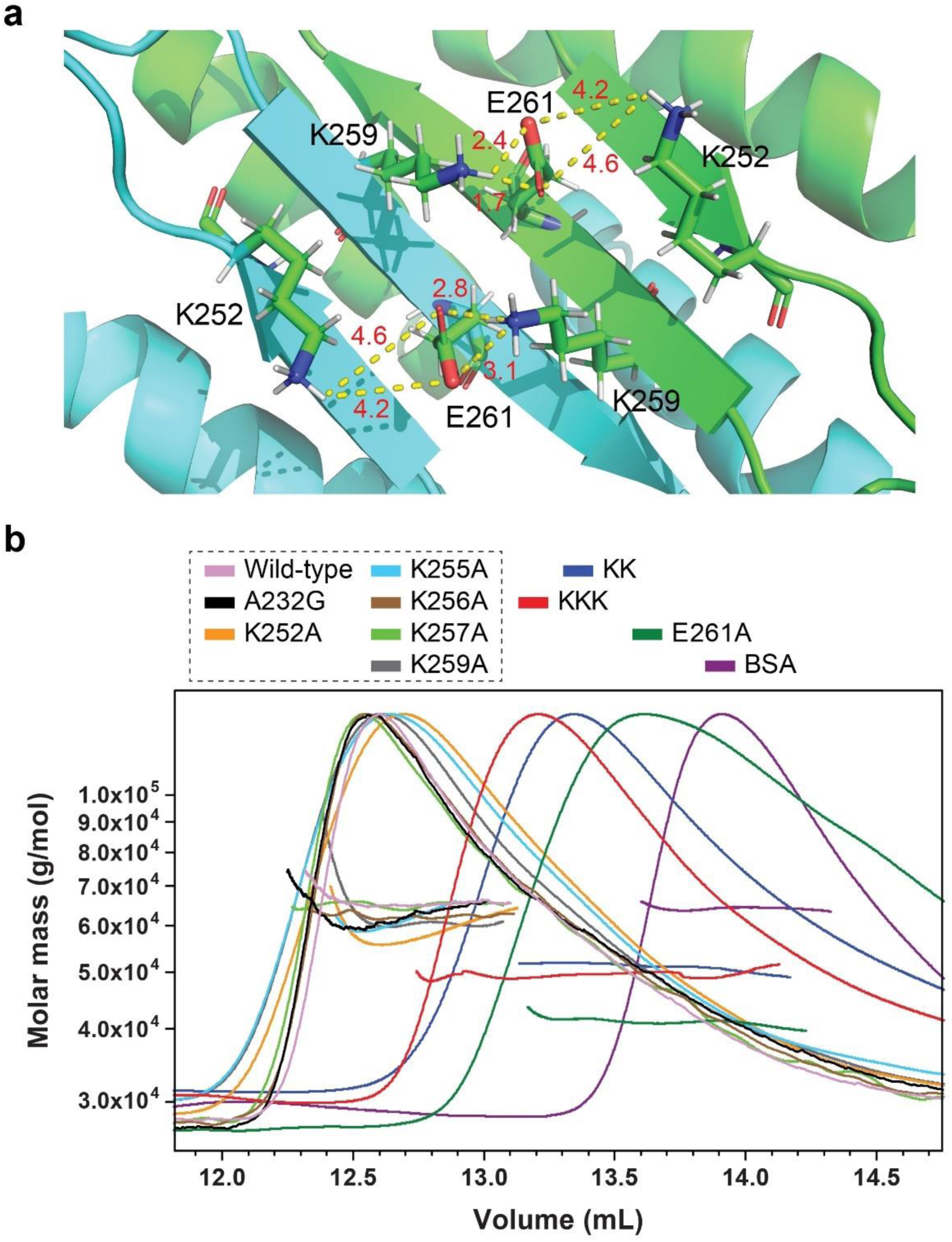
Salt bridge disruptions substantially weaken dimerization of the C-terminal domain. **a**, PyMol representation of salt bridge network formed by K252, K259, and E261 (PDB: 5NOC). Green and cyan colors represent each C-terminal domain (CTD) monomer. Numbers in red are distances in Angstroms. **b**, SEC-MALS (size exclusion chromatography coupled with multi-angle light scattering) characterization of wild-type and mutant ParB proteins. Light scattering signals are represented in the Rayleigh ratio. Signals were monitored using SEC-MALS and normalized to the maximum in each curve. 0.1 mL of 0.02 mM samples was injected with 20 mM Tris, 350 mM NaCl running buffer at pH 8.0.

The intra- and inter-monomer salt bridges mediated by E261 raised the possibility that those salt bridges play an instrumental role in CTD dimerization. We hypothesized that mutating E261 into alanine would disrupt the dimeric status of BsParB proteins. To investigate the monomer-dimer status of BsParB proteins, we employed size exclusion chromatography coupled with multi-angle light scattering (SEC-MALS) (Fig. 4b). A control experiment with bovine serum albumin (BSA; 66.5 kDa) correctly provided the molecular weight (64.2 kDa with 0.4% uncertainty) of its monomeric form (the major peak). As was observed from other ParB studies (27, 28, 46), the (full-length) wild-type BsParB in solution was eluted as a single peak with a calculated molecular weight of 65.7 kDa (uncertainty: 1.2%), which matches a theoretical dimeric molecular weight. However, the SEC-MALS measurements with BsParB(E261A) yielded a molecular weight of 41.1 kDa (with 2.0% uncertainty), which is clearly lower than the dimer‘s molecular weight (∼64 kDa) and slightly higher than the monomer’s (∼32 kDa). These results imply that BsParB(E261A) is primarily a monomer. While it runs through the size exclusion column, the two monomers can weakly and temporarily associate with each other. Our MALS system measures the average molecular weight of the monomer (major) and weakly associated dimer (minor) populations. The measured molecular weight of BsParB(E261A) proves our hypothesis that E261 is a key residue involved in the CTD dimerization. On the other hand, mutations on the lysine residues (K252A, K255A, K256A, K257A, and K259A) did not compromise the dimeric status of BsParB (59.0 kDa, 61.3 kDa, 62.4 kDa, 65.2 kDa, and 62.5 kDa with 2.4%, 2.2%, 1.5%, 1.6%, and 6.1% uncertainties, respectively) (Fig. 4b). The SEC-MALS results imply importance of the intra- and inter-monomer salt bridges mediated by E261 for CTD dimerization and raise questions about their roles in protein function.

### Molecular dynamics simulations of apo and DNA-bound CTD reveal conformational changes in the protein upon DNA binding

The absence of crystal or NMR structures for the DNA-bound CTD prompted us to perform molecular dynamics (MD) simulations of apo and DNA-bound CTD using GROMACS software. After equilibration procedures (Extended Data Fig. 4a-f), the MD simulation determined the order parameter S^2^ values (Fig. 3c,d). These values were compared with the experimentally obtained S^2^ values. In addition, we obtained and compared the average structures for the simulated apo- and DNA-bound CTD (Fig. 5a-d).

**Fig. 5.**
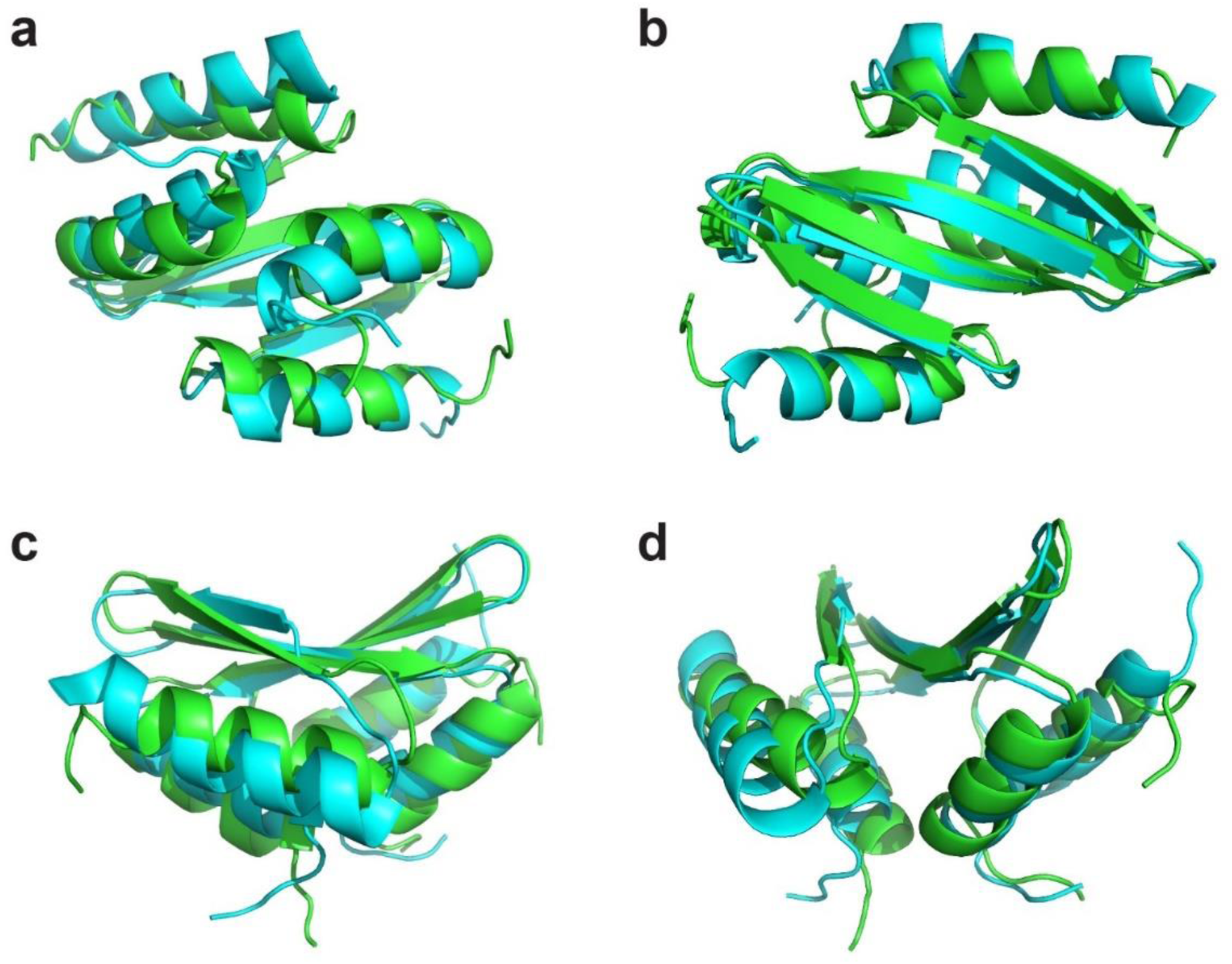
An overlay of average simulated apo CTD (green) and DNA-bound CTD (cyan) structures. Views from (**a**) the bottom, (**b**) the top, (**c**) the side, and (**d**) the front.

During 500 ns of simulation under identical conditions, both apo and DNA-bound CTD preserved their structural stability. We used variable averaging times (100 ps to 5 ns) to extract S^2^ order parameters from the MD trajectories (Supplementary Tables 6 and 7). The S^2^ values derived from MD for both the apo and the DNA-bound CTD are generally close to agreement with experimentally derived S^2^ (Extended Data Fig. 5a,b): (1) elevated mobility (lower S^2^) within the linker area (residues 253-257) and the termini, and (2) elevated rigidity (higher S^2^) within the rest of the backbone chain. The MD-derived S^2^ values for the apo CTD correctly predict K257 as the most flexible lysine residue outside of the termini regions, corroborating the NMR relaxation results. However, we noted that MD exaggerates internal motion (low S^2^) at the linker region, particularly in the apo CTD (Extended Data Fig. 5a,b). The exaggerations of NMR S^2^ values for flexible protein residues are a well-known feature of MD simulations (47, 48).

The general patterns of DNA binding effects on the S^2^ values were approximately reproduced in the MD simulations, except for the C-terminal half of α-helix 2 and the C-termini (Fig. 3e-f). First, positive “S^2^(DNA-bound)-S^2^(apo)” values in the middle of α-helix 1 and negative values from both ends of α-helix 1 were also obtained in the simulations, although the magnitude of the S^2^ value changes in the MD simulation was less pronounced. Consistent with experimental data, increased flexibility (lower S^2^) on β-helix 1 upon DNA-binding was observed, and the internal backbone rigidities for K256-K257 were increased upon DNA binding. However, contrary to the experimental data, DNA binding increased the K255 backbone rigidity in the simulation.

The comparison of the simulated CTD structures in the absence and presence of DNA (Fig. 5a-d) shows that α-helices of DNA-bound CTD are out of the center of symmetry. Also, while the β-strands of both the DNA-bound and the apo-CTD are closely overlaid in the simulation, the first β-strand of the DNA-bound CTD is shorter, extending the coil-like linker regions. Interestingly, compared with the NMR structure of CTD (PDB: 5NOC), the first β-strand in both the apo and DNA-bound simulated structures appears longer (Extended Data Fig. 6). The second β-strands in 5NOC and DNA-bound simulated structures exhibit the same length but are similarly shorter than the simulated apo structure (Extended Data Fig. 6). We argue that the shortening of β-strands of DNA-bound CTD in MD simulations is probably due to the critical importance of decreased structural constraint for DNA recognition and sliding. However, due to the lack of DNA-bound NMR or crystal structures of CTD or the full-length ParB, the results should be interpreted with caution by comparing only simulated apo and DNA-bound CTD structures. The results of MD simulations allowed us to “theoretically” observe the conformational changes upon DNA binding to the CTD and suggest that DNA modulates the structure of CTD by shortening the β-strands.

### Mutating individual CTD lysine residues reduces single-molecule DNA compaction rates

In NMR, chemical shift perturbation (CSP) is a powerful and sensitive tool in investigating structural changes upon ligand binding (49). Consistent with the previous results by Fisher *et al*. (29), our CSP results show that K252, K255, K256, and K259 lysine residues, along with A232 in α-helix 1 (231–246), experience high perturbation upon DNA binding (Extended Data Fig. 7). Among them, K256 shows the highest CSP (0.297), and K255 exhibits a marginally high CSP (0.083). In contrast, DNA binding does not yield a noticeable chemical shift perturbation on K257 in the loop region (253–257) (See reference (29) and Extended Data Fig. 7).

To understand how key CTD residues contribute to the overall function of the ParB protein, we employed a single-molecule DNA flow-stretching assay. Single-molecule DNA flow-stretching is a widely employed, versatile methodology in DNA-protein interaction studies (50). Its high sensitivity allows detection of even minor changes in proteins of interest by monitoring end-to-end length changes of flow-stretched DNA (51). In this assay, one end of the bacteriophage λ-DNA was tethered to a microfluidic sample chamber surface, and the other end was labeled with a fluorescent quantum dot. Applying laminar buffer flow leads to DNA stretching and delivers proteins to the flow-stretched DNAs. The bridging activities of DNA-ParB or ParB-ParB (28, 33, 38, 46, 52–54) result in end-to-end length changes of flow-stretched DNA, which are visualized by monitoring the quantum dot position over time.

The compaction rates of flow-stretched DNA of the full-length wild-type BsParB were compared with those of five (full-length) mutant ParB proteins. First, BsParB(A232G), BsParB(K252A), BsParB(K255A), BsParB(K256A), and BsParB(K259A) were tested, as residues participating in direct DNA binding or undergoing conformational changes upon DNA binding are expected to have large CSP values (49). Despite its low CSP, BsParB(K257A) was another mutant we tested due to its particularly high backbone flexibility (low S^2^). BsParB(E261A) was also chosen to investigate the effects of the disruptions of salt bridges between E261 and rigid K252/K259 residues. In our study, since our goal is to understand BsParB’s nonspecific DNA binding activity, we used bacteriophage λ-DNA, which lacks any *parS* sequences, as the DNA substrate.

As we previously reported (51), 50 nM wild-type BsParB robustly compacted flow-stretched DNA in the absence of CTP and showed substantial compaction rate decreases in the presence of CTP (Fig. 6a,b and Supplementary Table 8). The BsParB(A232G) mutant protein behaved similarly to the wild-type BsParB with a comparable compaction rate and slower compaction in the presence of CTP (Fig. 6b). However, regardless of CTP, *C. crescentus* ParB (CcParB), which lacks the CTD lysine patches (13, 14), did not compact DNA (Fig. 6b). We observed that single lysine mutant proteins at their 50 nM concentrations lost substantial DNA compaction capability. No compaction events were observed for BsParB(K252A), BsParB(K257A), and BsParB(K259A) regardless of CTP or for BsParB(K256A) in the presence of CTP (Fig. 6a,b). Even for 50 nM BsParB(K256A) without CTP, the compaction rate was 12-fold slower than that of wild-type BsParB. The 50 nM BsParB(E261A) protein, in which salt bridges are disrupted, also did not compact flow-stretched DNA (Fig. 6b).

**Fig. 6.**
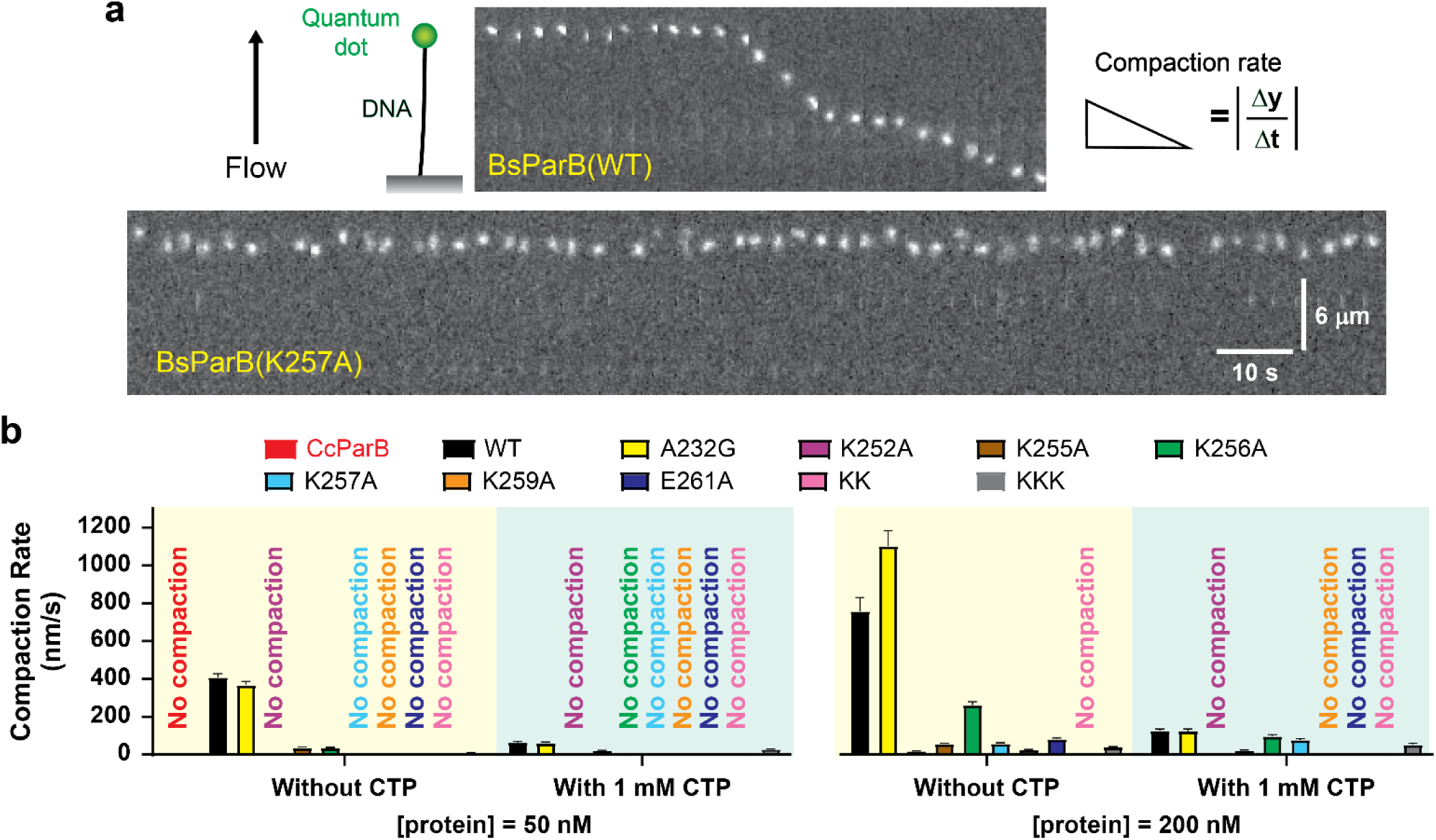
The CTD lysine mutations decrease the compaction rates of single-molecule flow-stretched DNAs. **a**, Schematic of single-molecule DNA flow-stretching assay. The two kymographs for the wild-type and the K257A mutant *Bacillus subtilis* ParB represent efficient and no DNA compaction events, respectively. **b**, DNA compaction rates by various ParB proteins (50 nM and 200 nM) both in the presence and absence of CTP (*n*=23∼234). CcParB stands for *Caulobacter crescentus* ParB, which lacks the C-terminal domain lysine patches. Other proteins are *B. subtilis* ParB. Error bars: SEM. Despite the presence of multiple lysines, a single lysine mutation results in a decrease in the DNA compaction rates.

Considering that multiple putative DNA-binding lysine residues exist at the CTD, we wondered whether a single-point mutation could abolish DNA compaction even at an increased protein concentration. DNA flow-stretching experiments with four-fold increased protein concentration (200 nM) showed that BsParB(K255A), BsParB(K256A), BsParB(K257A), and BsParB(K259A) mutants retained the DNA compaction abilities, albeit with significant disruption to DNA compaction (Fig. 6b and Supplementary Table 8). No DNA compaction was seen for 200 nM BsParB(K252A) with CTP. Our results demonstrate the crucial roles of CTD lysines in DNA compaction, most likely by being involved in DNA bridging events. Slow DNA compactions by 200 nM BsParB(E261A) were seen only when no nucleotides were supplemented. In summary, mutating individual lysines dramatically disrupts DNA compaction (despite that the remaining lysines are intact) (see Discussion later).

### K256A or K257A mutation does not affect fluorescent ParB foci formation *in vivo*

To investigate how ParB’s DNA compaction capability *in vitro* correlates with ParB’s cellular localization and foci formation *in vivo*, we performed live cell fluorescence imaging for *B. subtilis* strains harboring various mutant ParB proteins. GFP was tagged to the N-terminus of WT or mutant ParB, and the construct was expressed from an ectopic locus on the chromosome in which the endogenous *parB* gene was deleted (33, 51). Previous studies have shown that WT GFP-ParB form nucleoprotein complexes that appear as fluorescent foci (10, 11, 18, 29, 33, 35, 46, 51, 55–57), while BsParB(R80A) mutants exhibits diffusive localization (33, 46, 51). Fluorescent foci formation has been considered to represent *in vivo* ParB spreading in *B. subtilis* (33, 46). Consistent with previous work, we found that wild-type GFP-ParB showed clear foci, and GFP-ParB(R80A) failed to form foci (Fig. 7a). We used these two strains as a positive control and a negative control, respectively. Consistent with the *in vitro* results that BsParB(A232G) compacted DNA similar to the WT (Fig. 6b), fluorescent foci formations were similar for WT ParB and ParB(A232G) in cells (Fig. 7a). For the salt bridge-disrupted BsParB(E261A) mutant, fluorescence foci formation was in between those for WT GFP-ParB and GFP-ParB(R80A). Foci-like signals from GFP-BsParB(E261A) were still detected, but fuzzy and faint. Strikingly, bright GFP-BsParB(K256A) and GFP-BsParB(K257A) foci were observed *in vivo* (Fig. 7a), in contrast to the poor flow-stretched DNA compaction by BsParB(K256A) and BsParB(K257A) observed *in vitro* (Fig. 6b). These results suggest that for major defects in protein structure, *in vitro* DNA compaction assay and *in vivo* fluorescence foci formation show consistent results. However, *in vitro* DNA compaction assay is more sensitive than *in vivo* foci formation to reveal subtle protein structure changes. (See more in Discussion.).

**Fig. 7.**
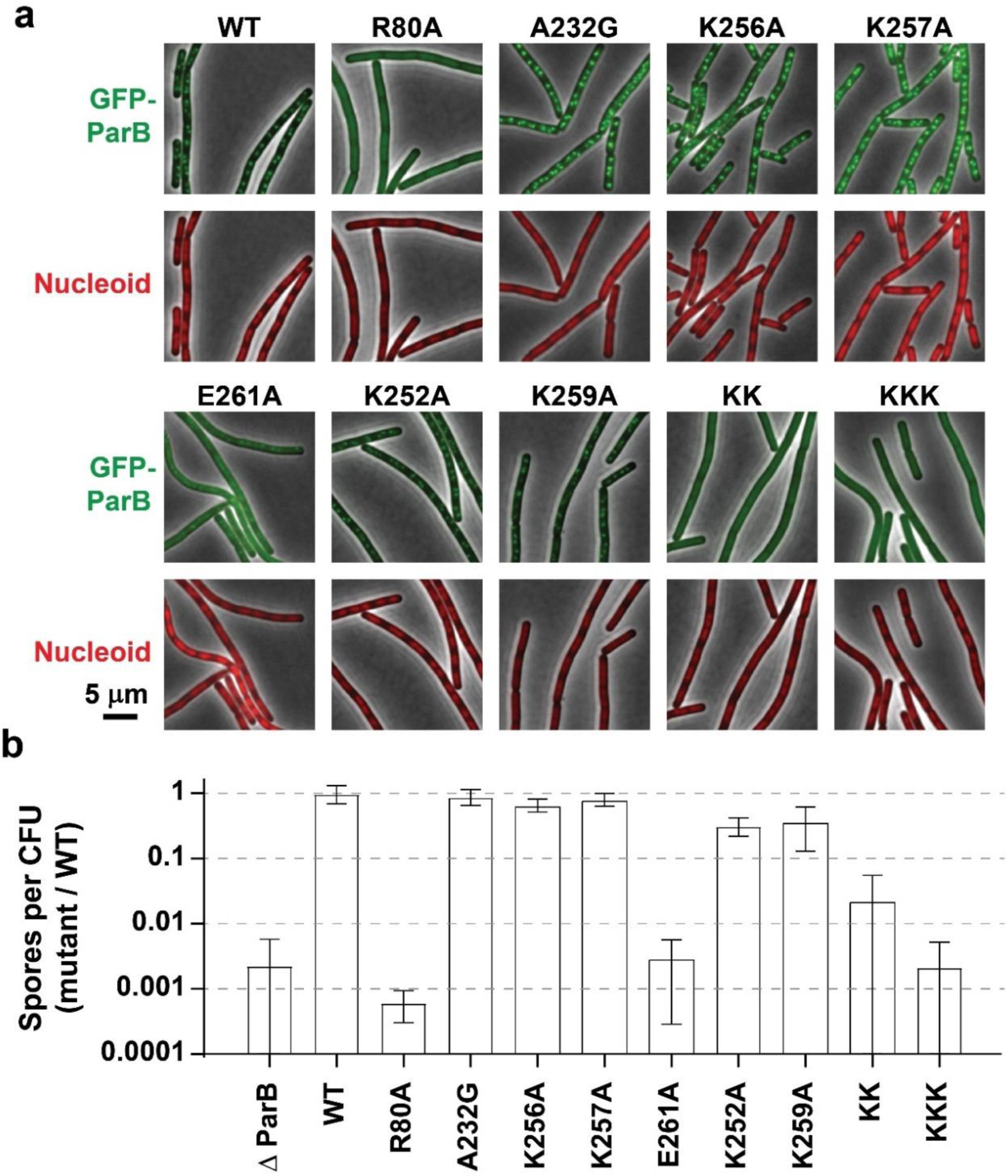
Disruption of CTD dimerization interferes with proper ParB foci formation and sporulation initiation. **a**, Visualization of GFP-ParB variants. WT and R80A serve as a positive control and a negative control, respectively. The E261A, “KK” (K252A, K259A), and “KKK” (K252A, K255A, K259A) mutations in GFP-ParB result in a loss of foci formation or the presence of faint foci. Cells were grown in CH medium and imaged during exponential growth using a Nikon Ti2 microscope. **b**, Sporulation efficiency (spore/CFU) relative to WT. Three biological replicates were averaged, and one standard deviation was represented by the error bars. Error bars extending to the x-axis (as shown for *ΔparB* and “KKK”) indicate that zero was included within the range of the standard deviation, which cannot be represented on a logarithmic scale.

### CTD dimerization is required for ParB’s *in vivo* activities

We noticed that the dimeric BsParB(A232G), BsParB(K256A), and BsParB(K257A) proteins form distinct *in vivo* nucleoprotein complexes like the wild-type counterpart, while dimerization-perturbed BsParB(E261A) entails noticeable foci disruptions. Based on these *in vivo* live cell imaging and SEC-MALS results, we hypothesized that ParB protein dimerization via the CTD is essential for ParB protein actions *in vivo*.

We first attempted to identify other CTD mutations that affect dimerization. Given that E261 forms salt bridges with K252 and K259, we purified BsParB(K252A-K259A) (hereafter “BsParB(KK)”) and BsParB(K252A-K255A-K259A) (hereafter “BsParB(KKK)”) to investigate their monomeric/dimeric statuses using SEC-MALS. We did not include the monomer candidate BsParB(L270D-L274D), where leucine zipper-mediated dimerization is abrogated, because it has been reported that this mutant is insoluble upon overexpression (29).

The molecular weights of BsParB(KK) and BsParB(KKK) were measured to be 51.2 kDa (with 2.0% uncertainty) and 49.5 kDa (with 1.4% uncertainty), respectively (Fig. 4b), much smaller than the molecular weight of the ParB dimer. These values suggest disruptions in CTD dimerization because dynamic monomer and dimer transitions result in intermediate molecular weight values in SEC-MALS experiments and elution as a single peak through the size-exclusion column. And, the DNA compaction abilities of those mutants were mostly lost (Fig. 6b). Next, we investigated how these mutations affect the formation of fluorescent ParB foci in live cells. While *B. subtilis* cells harboring dimerized BsParB(K252A) or BsParB(K259A) exhibit clear fluorescent foci, foci formation was abrogated in BsParB(KK) and BsParB(KKK), which is consistent with CTD dimerization disruptions (Fig. 7a).

Besides being important for chromosome segregation, ParB protein plays a critical role in initiating sporulation in *B. subtilis* (16), which is abolished by the R80A mutant (58). To investigate whether the CTD mutations mentioned above impact spore formation, we analyzed sporulation efficiency (Fig. 7b) . As a negative control, we measured the sporulation efficiency for *ΔparB*, which is 0.23±0.35% (mean±standard deviation) relative to the wild-type strain, consistent with previous results (∼1.2%) (16). Similar to *ΔparB*, ParB(R80A) had a relative sporulation efficiency of 0.06±0.03%. When the CTD mutations were introduced to endogenous *parB* locus and tested in the same way, the sporulation efficiencies of cells containing BsParB(A232G), BsParB(K256A), or BsParB(K257A) were comparable to the wild-type strain (89.0±24.5%, 66.4±14.9%, and 81.40±18.60%, respectively.) For cells containing BsParB(K252A) or BsParB(K259A), the sporulation efficiencies were slightly reduced, to approximately a third of the wild-type strain (Fig. 7b). Importantly, E261A (0.30±.0.27%), K252A-K259A (KK; 2.29±3.21%), and K252A-K255A-K259A (KKK; 0.22±0.30%) mutations reduced spore formation to a level similar to *ΔparB*.

Therefore, from SEC-MALS, *in vivo* live cell imaging, and sporulation assay, we discovered a prevalent trend: disruption of the ParB CTD dimerization, as seen from BsParB(E261A), BsParB(KK), and BsParB(KKK), is directly correlated with *in vivo* ParB activities seen in foci formation and sporulation initiation. Thus, the dimerization of the CTD is required for ParB’s *in vivo* functions.

## Discussion

Thermal fluctuations of structures allow proteins to continuously explore different conformational substates (39, 59). It has been shown that protein backbone flexibility on the picosecond-nanosecond (ps-ns) timescale facilitates DNA-binding proteins in navigating nonspecific DNA (60). In this study, we aimed to understand how these fast backbone dynamics (in ps-ns time scale) of the lysine-rich DNA-binding CTD surface are translated into functions of ParB using various *in vitro*, *in vivo*, and molecular dynamics simulation approaches.

First, we used NMR relaxation approaches which reveal ps-ns protein backbone dynamics with atomic resolution (59). We show that the R2/R1 ratio along the CTD backbone is higher for the N-terminal half than for the C-terminal half (Fig. 1c,g), suggesting elevated conformational dynamics of secondary structures on the C-terminal side of the CTD in comparison to the N-terminal side. These results indicate that CTD has asymmetric structural and functional differences along its backbone. Interestingly, in the presence of DNA, the number of residues affected by the additional conformational exchange was reduced from twelve to three, suggesting that DNA stabilizes the CTD structure. Upon DNA binding, NOE increases for secondary structures on the N-terminal side and on the linker but decreases for secondary structures on the C-terminal side of CTD (Fig. 1d,h), indicating that DNA binding elevates conformational dynamics within the C-terminal side.

The experimental data obtained from the NMR measurements were used to calculate the simulated order parameters (S^2^) for residues in the CTD. We found that DNA-binding lysine residues exhibit different dynamics. The lysine residues on the linker region (253–257) show significant flexibility, while those on β-strands (K252 and K259) show high rigidity (high S^2^). The rigid K252 and K259 not only participate in protein-DNA interaction, as we can observe it from CSP data, but also in the dimerization of CTD by forming salt bridges with E261. These findings obtained from the experimental NMR are supported by our molecular dynamics simulations, both showing that DNA-protein interface is formed by rigid and flexible regions. DNA slides along the rigid region with help from flexible lysine residues. These results show the importance of protein local dynamics in its function.

We found that single lysine mutations in ParB CTD (K252A, K255A, K256A, K257A, or K259A) exhibited substantially decreased DNA compaction rates in single-molecule DNA flow-stretching experiments, indicating that these lysine residues participate in nonspecific DNA binding and are important for *in vitro* DNA compaction. Consistent with this idea, *C. crescentus* ParB (CcParB), which lacks positively charged lysine patches at its CTD (13, 14), is incapable of compacting DNA *in vitro* (Fig. 6b), unlike ParBs from P1 plasmid, *Streptococcus pneumoniae*, *Pseudomonas aeruginosa*, *B. subtilis* or ParB1 from *Vibrio cholerae,* which contain the lysine patches and are capable of compacting flow-stretched DNAs (33). In the past, for BsParB(R80A) and many other *B. subtilis* mutants, the *in vitro* DNA compaction rates in the single-molecule DNA flow-stretching experiments were found to be coupled with *in vivo* fluorescence foci formation and sporulation (16, 33). Surprisingly, single lysine mutations (K252A, K256A, K257A, or K259A) did not hamper the *in vivo* functions of *B. subtilis* cells as analyzed by fluorescence foci formation in live cells and sporulation efficiency. The weak correlation between *in vitro* DNA bridging and *in vivo* functions has also been reported in other mutants in a separate study (46). Specifically, BsParB(M104A) and BsParB(Q140A) had reduced capability to compact DNA *in vitro* but formed fluorescence foci like the wild-type *in vivo* (46). The inconsistency implies that the *in vitro* single-molecule DNA compaction assay is more sensitive at detecting changes in protein structure than *in vivo* assays.

Our results raise a question of how ParB’s *in vivo* functions tolerate defects in interactions between DNA and ParB CTD. In *B. subtilis*, spore formation involves both ParA (Soj) and ParB (Spo0J) proteins (61). Null mutations in *parB* lead to an 80-200 fold decrease in sporulation initiation (Fig. 7b and (16)) compared with the wild-type strain. Importantly, the *parA* null mutation suppressed the sporulation initiation defects caused by *ΔparB* (9). Thus, it is the misregulation of ParA in Δ*parB* that causes sporulation defects (16, 61). It has been shown that it is ParB’s NTD that interacts with ParA and stimulates ParA ATPase (1, 13, 14). Thus, single lysine mutations within the CTD (K252A, K256A, K257A, or K259A) still allow ParB-ParA interactions. For *in vivo* fluorescence foci formation, despite the presence of nine *parS* sites on the *B. subtilis* chromosome (4), the wild-type GFP-ParB proteins appear as a single focus under fluorescence microscopy. Marston and Errington discovered that the absence of ParA leads to multiple smaller fluorescent ParB foci (62), suggesting that ParA proteins condense ParB-*parS* nucleoprotein complexes into one compact cluster (61, 62). Our results are consistent with this idea, suggesting that *in vivo* ParA helps single CTD (dimeric) lysine mutants form fluorescent foci. Similarly, our previous study showed that adding a small KCK- or ECE-tag to ParB protein changes *in vitro* DNA compaction rates but does not alter *in vivo* foci formation or ParB distribution along the chromosome (51), which might also be explained by the help of ParA. Finally, it is also possible that other cellular factors, like molecular crowding and DNA supercoiling, allow mild DNA-binding defects to be tolerated. Future studies are needed to validate these possibilities.

Our *in vitro* single-molecule DNA flow stretching results (Fig. 6) and live cell imaging and sporulation assays (Fig. 7a,b) showed that BsParB(E261A), BsParB(KK), and BsParB(KKK) exhibit strong defects in DNA compaction and *in vivo* functions. Those three mutants exhibit reduced CTD dimerization as judged by SEC-MALS (Fig. 4b). These results show that ParB dimerization via CTD is essential for *in vivo* ParB function and support the ParB sliding clamp model (19, 20, 26, 36, 37) where DNA is trapped in the compartment between the DNA-gate and the CTD.

The ParB sliding clamp model is reminiscent of the structure and function of proliferating cell nuclear antigen (PCNA), the DNA sliding clamp in eukaryotic cells. A ring-shaped homotrimer PCNA encircles DNA and plays a crucial role in DNA replication and damage repair (64). Crystal structures of PCNA revealed that a central channel (inner rim) is lined with positively charged lysines and arginines to interact with the negatively charged DNA backbone (Fig. 8a) (65, 66), similar to the arrangement of lysine side chains on the BsParB CTD. The basic (lysines and arginines) PCNA residues match the B-form DNA helix pitches. Short-lived polar interactions (on a sub-nanosecond timescale) between PCNA basic residues and DNA were proposed to facilitate PCNA sliding on DNA (67). Inspired by the PCNA inner rim geometry, we inspected the DNA-BsParB CTD interface using the CTD NMR structure (5NOC). We observe that the distances between lysines forming the DNA-BsParB CTD interface are close to the dimensions of the major and minor grooves of DNA (Fig. 8b). The distance between the flexible lysines on the edge of the interface (K255-K257) and the lysines at the middle (K252 and K259) is approximately 14 Å, which fits inside the major groove of DNA (around 22 Å) (68). The dimensions formed by four lysines at the center of the DNA-BsParB CTD interfaces (K252 and K259 from each monomer) are between 6 Å and 7 Å, which fit inside the minor groove (around 12 Å) (68). We propose that geometries formed by lysines at the DNA-CTD interface promote DNA sliding along the surface of the ParB CTD. Lastly, despite multiple DNA-interacting basic residues at the PCNA inner rim, mutating individual lysines or arginines to alanine leads to disruptions of PCNA-associated functions, such as the processivity of polymerase δ and the movement of PCNA on DNA (69, 70). Those impacts were ascribed to the disruptions of clamp loading and the improper positioning of DNA along the basic PCNA inner rim residues (70). Intriguingly, mutating only one BsParB CTD lysine residue substantially lowered the single-molecule DNA compaction rates (Fig. 6b). The origin of the dramatic effects from a single point mutation is elusive, but the predicted DNA-BsParB CTD AlphaFold 3 (71) structure, where DNA is aligned along the CTD lysine residues, provides a hint (Fig. 8c). As proposed in PCNA, we speculate that the BsParB CTD lysine residues collectively position a nonspecific DNA, so that the disruption of any of these lysine residues misaligns the DNA. Overall, our NMR S^2^ and single-molecule results, along with inspiration from PCNA, suggest a “rail” mode of DNA non-specific binding where rigid (high S^2^) K252 and K259 form an “inner guide,” while flexible (low S^2^) outer K255-K257 promote efficient DNA sliding.

**Figure 8.**
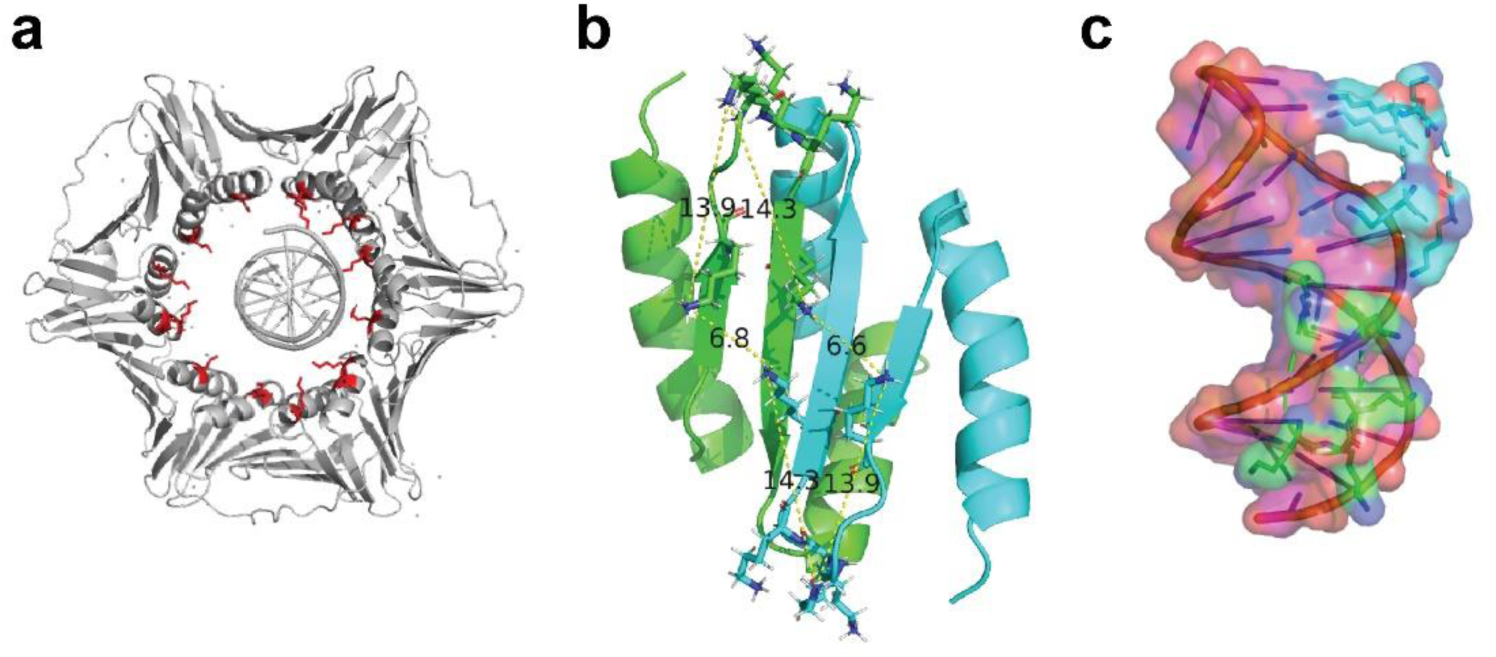
The configuration of the ParB CTD lysines enables the proper positioning of DNA. **a**, A structure of human proliferating cell nuclear antigen (PCNA) and associated DNA (PDB: 6GIS). The inner rim basic (lysines and arginine) residues are shown in red sticks. The arrangements and fast dynamics of these basic residues enable efficient DNA sliding. **b**, The distances between lysine residues (stick representation) on *B. subtilis* ParB CTD (PDB: 5NOC) are shown in angstroms. Green and cyan colors represent different monomer CTDs. **c**, AlphaFold 3 structure of DNA-bound ParB CTD. For clarity, only lysine residues on each monomer are shown in green and cyan, and the rest of the ParB CTD is omitted. The ParB CTD Lysines are aligned along the DNA backbone.

In summary, our NMR relaxation and molecular dynamics studies reveal that CTD lysines exhibit distinct dynamics, facilitating non-specific DNA binding to enable efficient ParB sliding along the DNA. Single-molecule approaches and AlphaFold 3-based structural prediction suggest that the integrity of all lysine residues is crucial for nonspecific DNA binding. Any single lysine mutation significantly reduces DNA binding of ParB CTD. However, single lysine mutations do not affect ParB’s *in vivo* function, suggesting that other cellular factors, such as ParA-ParB interactions via NTD, molecular crowding, and DNA supercoiling, play a role. Only when ParB dimerization via the CTD is compromised (like in BsParB(E261A), BsParB(KK), and BsParB(KKK)), *in* vivo ParB functions are disrupted. Our results are consistent with the formation of ParB clamp after loading on *parS*. Our work sheds light on mechanisms of ParB and reveal the impact of rapid protein dynamics on protein functions. Lastly, the remarkable similarities in the fast dynamics of ParB and PCNA raise questions about whether other DNA-binding proteins that move on DNA, such as DNA translocases and proteins involved in homology searches, adopt similar strategies.

## Acknowledgements

Support for this work comes from National Institutes of Health R01GM141242, R01GM143182, R01AI172822 (X.W.), and R35GM143093 (H.K.). This research is a contribution of the GEMS Biology Integration Institute, funded by the National Science Foundation DBI Biology Integration Institutes Program, Award #2022049 (X.W.).

## Methods

### Plasmids and DNA preparation

The plasmids with the coding sequence of His6-SUMO-tagged *B. subtilis* ParB (BsParB) proteins were constructed as previously described by assembling double-stranded gBlocks^TM^ gene fragments. Briefly, a polymerase chain reaction was performed with the plasmid that encodes for His6-SUMO-BsParB(wild-type) (m0041 = pTG011) to generate a linearized vector devoid of a part of the SUMO-tag coding sequence and the entire wild-type BsParB. Double-stranded gBlocks^TM^ gene fragments in which sequences encode for the C-terminal domain or mutant BsParB proteins were purchased from Integrated DNA Technologies (Coralville, IA). The desired plasmids were generated from the linearized vector and synthesized inserts by following the NEBuilder HiFi DNA Assembly Master Mix (NEB E2621S, Ipswich, MA) protocol. The reaction mixtures were used to transform NEB 5-alpha competent *E. coli* cells. After performing minipreps, Sanger sequencing (Psomagen, Rockville, MD) was used to confirm the sequences of the plasmids.

Ten bp DNA hairpins for NMR relaxation studies were prepared as previously reported by Fisher *et al*. (29) and concentrated in phosphate-buffered saline (PBS) buffer (pH 6.1).

### Culture growth and *B. subtilis* ParB C-terminal domain purification for NMR study

The plasmid encoding the His6-SUMO-BsParB(217–282) construct (m0083) was transformed into BL21(DE3)pLysS *Escherichia coli* competent cells. 3 mL of Luria-Bertani (LB) broth was inoculated with the transformed cells from a single colony in the presence of 100 μg/mL ampicillin and 20 μg/mL of chloramphenicol. After growing the cells for 5-6 hours at 37°C in a shaking incubator (250 rpm), 0.8 mL of the cell culture was transferred into 40 mL of 90:10 (vol/vol) M9 minimal/LB media that contains U-^15^N NH4Cl and incubated overnight at 37°C. The next morning, 20 mL of the overnight culture was transferred into 1 L of M9 minimal media and allowed to grow at 37°C while shaking at 250 rpm. Once the OD600 reached 0.6-0.8, protein overexpression was induced by adding isopropyl β-D-1-thiogalactopyranoside (IPTG; 0.5 mM final concentration). After induction by IPTG, the cell culture was allowed to grow at 37°C for 4 hours. Cells were harvested by centrifugation. The cell pellets were resuspended in lysis buffer (20 mM Tris, pH 8.0, 1 M NaCl, 20 mM Imidazole) containing a Roche protease inhibitor cocktail (Roche, Basel, Switzerland) and 0.1 mM phenylmethylsulfonyl fluoride (PMSF) and then flash-frozen.

His6-SUMO-BsParB(217–282) (BsParB C-terminal domain) protein was purified following the previously reported protocol (33, 46, 72). After thawing the cells, we added additional PMSF to 1.0 mM final concentration, 0.25 mg/mL lysozyme, 750U universal nuclease (Thermo Fisher Scientific 88701), and 5.0 mM 2-mercaptoethanol (βME). After 30 min on ice, cells were sonicated with a microtip (amplitude 40%, total time 15 min, pulse on: 4.0 sec, pulse off: 2.0 sec) on Branson SFX150 sonifier. The cell lysate was first centrifuged at 11,000 g with an FA-6x50 rotor in the Eppendorf 5910R centrifuge for 30 min at 4°C. The supernatant was once again centrifuged at 20,133 g for 30 min. The supernatant containing His6-SUMO-BsParB(217–282) was collected and mixed with 6 mL of Ni-NTA agarose solution (Qiagen #30230) for a 3 mL bed volume and supplemented with one tablet of Complete Mini-EDTA-free protease inhibitor cocktail (Roche 04693159001). The supernatant was incubated for 1 hour. After applying the mixture of supernatant and Ni-NTA resin, we washed the column with 30 mL of Lysis buffer supplemented with 5 mM MgCl2 and 5 mM βME. Then, we washed the column with salt reduction buffer (20 mM Tris, pH 8.0, 350 mM NaCl, 20 mM Imidazole, 5 mM MgCl2, 5 mM βME). Finally, proteins were eluted with Elution buffer (20 mM Tris, pH 8.0, 350 mM NaCl, 250 mM Imidazole, 5 mM MgCl2, 5 mM βME). After each step, we collected fractions and analyzed them using an SDS-polyacrylamide (SDS-PAGE) gel. Fractions with identified His-SUMO-BsParB(217–282) were combined, supplemented with SUMO cleaving His-tagged Ulp1 protein, and left to incubate for 1 hour at 4°C. After 1 hour, the protein solution was transferred into a dialysis tube. We dialyzed the protein three times (1 hour in 500 mL, overnight in 1 L, and 2.5 hours in 500 mL) against ParB dialysis/storage 1 buffer (20 mM Tris, pH 8.0, 350 mM NaCl, 10 mM imidazole, 5 mM βME, 1 mM MgCl2, 10% glycerol). Collected from the dialysis bag, the protein solution is remixed with the Ni-NTA resin and incubated for 1 hour. The mixture was added to a 5 mL polypropylene column (Qiagen), and the unbound CTD (devoid of His-SUMO) protein (BsParB(217–282)) was eluted. ParB dialysis/storage 2 buffer (20 mM Tris, pH 8.0, 350 mM NaCl, 10% glycerol) was used to elute more unbound CTD. The collected fractions were analyzed with an SDS-PAGE gel. Pure CTD fractions were consolidated. The final protein was snap-frozen in liquid nitrogen and stored at -80°C. CTD concentration was determined by UV-Vis using theoretical extinction coefficients of 2560 M^-1^ cm^-1^.

### Overexpression and purification of full-length *B. subtilis* ParB proteins

Full-length ParB proteins were overexpressed and purified based on the overexpression and purification protocol for His6-SUMO-BsParB(217–282) protein except for the following modifications: (1) Rosetta2(DE3)pLysS cells were transformed, and only LB medium was used in culturing the transformed cells. (2) After adding IPTG, the culture was grown for 4 hours at 30°C. (3) the sonication was done at 30% amplitude, for a total of 10 seconds with 1 second pulses on and 1 second pulses off. (4) Apyrase was supplemented into the clarified supernatant to deplete cellular NTPs. (5) For the protein concentration determination, 7450 M^-1^ cm^-1^ was used as the theoretical extinction coefficient.

### NMR sample preparations

Uniformly ^15^N-labeled samples for solution NMR measurements were prepared from purified protein by buffer exchange in phosphate-buffered saline (PBS) (1.8 mM KH2PO4, 10 mM NaH2PO4, 137 mM NaCl and 2.7 mM KCl (pH 6.1)), with 10% D2O and 200 mM 4,4-dimethyl-4-silapentane-1-sulfonic acid (DSS). NMR samples were prepared at 0.4 mM protein concentration and loaded into 1.3 mm NMR tubes. DNA-bound protein samples were prepared by adding the ten-base pair hairpin DNA (GCGTACATCATTCCCTGATGTACGC) in 0.25 to 1.25 equivalents.

NMR datasets for ^1^H-^15^N HSQC (Heteronuclear Single Quantum Coherence) (73), ^1^H-^15^N T1 (74, 75), T2 (75), and heteronuclear NOE (Nuclear Overhauser Effect) bi-dimensional experiments (76) were collected at 308 K, utilizing a Bruker NMR 500 MHz spectrometer with a 1.7 mm cryogenic cold-probe located at UT Health of San Antonio.

### 15N Relaxation Experiments

We used the two-dimensional ^1^H-^15^N HSQC pulse sequence “hsqct1etf3gpsi3d” to measure heteroatom T1 relaxation time (40, 77). A pseudo three-dimensional ^1^H-^15^N HSQC spectra for longitudinal relaxation time T1 of CTD was acquired in Q.F. mode for proton dimension (F1) and Echo-Antiecho mode for nitrogen dimension (F2). The spectral widths were 16 ppm for direct and 35 ppm for indirect dimensions, respectively. Proton and nitrogen carrier frequencies were centered at 4.703 ppm and 117 ppm, respectively. The time points collected were 2048 for the proton dimension and 256 for the nitrogen dimension. The acquisition time was 0.000564 seconds for proton and 0.0722 seconds for nitrogen. The number of scans acquired was 32. ^15^N T1 values were measured from the spectra and recorded with seven different durations of the delay T: T = 0.01, 0.08, 0.2, 0.4, 0.6, 0.8, 1.0 seconds. The measurements were repeated three times for a delay of 0.2 seconds. The experiment was recorded for one day and eight hours. We used the two-dimensional ^1^H-^15^N HSQC pulse sequence “hsqct2etf3gpsi3d” to measure heteroatom T2 relaxation time (78). A pseudo three-dimensional ^1^H-^15^N HSQC spectra for transverse relaxation time T2 of CTD were acquired in Q.F. mode for proton dimension (F1) and in Echo-Antiecho mode for nitrogen dimension (F2). Proton and nitrogen carrier frequencies were centered at 4.703 ppm and 117 ppm, respectively. The spectral width was 16 ppm for direct and 35 ppm for indirect dimensions. The time points collected were 2048 for the proton dimension and 256 for the nitrogen dimension. The acquisition time was 0.000689 seconds for the proton dimension and 0.0722 seconds for the nitrogen dimension. The number of scans acquired was 32. ^15^N T2 values were measured from the spectra recorded with nine different durations of the delay T: T = 0, 0.016, 0.032, 0.048, 0.064, 0.080, 0.096, 0.112, and 0.128 seconds. The measurements were repeated three times for a delay of 0.032 seconds. The experiment was recorded for one day and eight hours.

For the heteronuclear NOE pulse sequence “hsqcnoef3gpsi3d” (79), spectra were recorded in the presence and absence of ^1^H saturation: 4 s saturation plus 1 s recycle delay, and the reference experiment with 5 s recycle delay. Heteronuclear ^15^N-^1^H NOE spectra for CTD were acquired in QF (no frequency) mode for the proton dimension (F1) and in Echo-Antiecho mode for the nitrogen dimension (F2). Proton and nitrogen carrier frequencies were centered at 4.703 ppm, and 117 ppm, respectively. The spectral width was 16 ppm for direct and 35 ppm for indirect dimensions. The time points collected were 2048 for the proton dimension and 512 for the nitrogen dimension. The acquisition time was 0.0002 seconds for the proton dimension and 0.144 seconds for the nitrogen dimension. The number of scans acquired was 32. The experiment was recorded for one day and fifteen hours.

### NMR Data Processing

All obtained NMR spectra were processed with NMRpipe (80) and analyzed with Sparky (version 3.113. The University of California, San Francisco). A forward linear prediction to twice the number of original data points followed by zero filling to twice the total number of points was used for all experiments prior to Fourier transformation. 45°-shifted sine bell apodization with a water suppression function was used for both dimensions.

### Determination of R1, R2, NOE and Modelfree Calculations

To determine R1 and R2, the T1, T2, and ^15^N{^1^H} NOE relaxation data obtained by NMR relaxation experiments were analyzed with the routine available within SPARKY. We measured the intensities of the ^15^N-^1^H cross-peaks from the relaxation experiments for the various recovery delay times using SPARKY software. The peak intensities for the protein residues were then fitted to the exponential decay equation (1) by the SPARKY routine to determine T1 or T2, where T1 and T2 are the relaxation rates. Knowing T1 or T2, we calculated R1 and R2 as R1= 1/T1 and R2 = 1/T2.

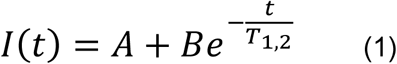

The errors for both R1 and R2 were calculated from estimated T1 and T2 errors from the fitting routine. The {^1^H}-^15^N Heteronuclear NOEs were calculated from the ratio of cross-peak intensities in the two experiments: with and without proton saturation.

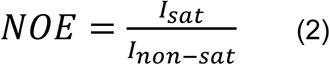

r2r1_tm and quadric_diffusion routines developed by the Palmer group (81) were used to determine the initial value for overall correlation time (τm) and the initial estimate of the diffusion tensor. The latter was a prerequisite to determine the appropriate motional model (spherical, axially symmetric, or fully anisotropic). r2r1 routine calculates R2/R1 ratio of the individual residues ^15^N amide peaks. The calculated τm values are then fed into the quadric_diffusion routine. To calculate the initial estimate of the diffusion tensor, along with the τm values, we used the PDB file of the NMR structure of apo CTD (PDB entry 5NOC) modified by the “pdbinertia” program also developed by the Palmer group (81).

To select the dynamic model describing the residues’ internal motion, we used the FAST-ModelFree program developed by Loria and Cole (82). This program performs model-free calculations based on model-free formalism developed by Lipari and Szabo (44). In short, model-free formalism states that if molecular global tumbling and internal motions have enough separation (at least one order of magnitude), these processes can be decoupled. The model-free formalism allows the data from relaxation experiments (R1, R2, NOE) to be analyzed by fitting several “model-free” spectral density functions (Supplementary Table 2). Through these calculations, we can obtain two essential parameters: S^2^ and τ. S^2^ is an order parameter reflecting the rigidity of the vector, N-H bond in the case of protein, with τ being a timescale of the vector’s internal motion. The FAST-Modelfree conducts the model calculation through the Modelfree program written by the Palmer group (39, 83, 84), which fits a model function for each residue. Our input file for the FAST-Modelfree program had an N−H bond length set to 1.02 Å and ^15^N chemical shift anisotropy (CSA) equal to 172 ppm.

### Chemical shift perturbation analysis

The following equation calculates chemical shift perturbations (CSP) for backbone amides between initial and final states 1 and 2 (39).

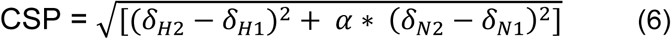

*δ*_*H*1,2_ and *δ*_*N*1,2_ values are the ^1^H and ^15^N chemical shifts for states 1 and 2, respectively, with α = 0.2.

### DNA substrate preparation for single-molecule experiments

The overhangs of bacteriophage lambda DNA were employed to tag one end with biotin and the other end with digoxigenin, as outlined in our previous works (85). This allows lambda DNA tethering to the microfluidic flow cell surface through neutravidin-biotin interactions facilitated by biotin. The opposite end of lambda DNA is labeled with digoxigenin, essential for attaching anti-digoxigenin antibody-conjugated quantum dot 605 (Invitrogen, Waltham, MA). Briefly, a biotinylated oligo was annealed to a complementary overhang, followed by ligation, and a similar process was carried out for digoxigenin-oligo. The removal of excess short oligos was executed through electrophoresis, and DNA substrates were obtained via ethanol precipitation.

### Microfluidic flow cell preparation

Cover glasses were effectively surface-passivated using (3-aminopropyl) triethoxysilane (Millipore Sigma A3648, St. Louis, MO) followed by a mixture of polyethylene glycol (PEG; MPEG-SVA-5000-1g) and its biotinylated version (Biotin-PEG-SVA-5000-100mg) (Laysan Bio, Arab, AL) as stated in previous publications (85, 86). A microfluidic flow cell was constructed by drilling parallel holes on a quartz plate (Technical Glass Product, Paineville, OH) and placing double-sided tape (Grace Bio-Labs, Bend, OR) between a PEGylated cover glass and the modified quartz plate. The parallel-affixed double-sided tapes create a rectangular cell channel with a defined height and width. The inlet tube (7 cm) and outlet tube (2.5 cm) were inserted through the holes in the quartz plate and sealed air-tight using epoxy. The PE60 inlet tube at one channel end was immersed in a buffer-containing tube, while the outlet tubes at the other end were connected to a syringe on a syringe pump (Harvard Apparatus, Holliston, MA).

### Single-molecule DNA flow-stretching and data analysis

Approximately 4% of the PEG on the surface-passivated cover glass contains biotins, acting as a neutravidin binding platform. 0.25 mg/mL of neutravidin was administered, followed by quantum dot-labeled biotinylated lambda DNAs. The removal of unlabeled quantum dots and untethered DNAs was achieved by washing the flow cell with an imaging buffer (10 mM Tris, pH 8.0, 150 mM NaCl, and 10 mM MgCl2). Movie acquisition was acquired after a minimum of two minutes without buffer flow to ensure that the average quantum dot position coincides with the DNA tether point due to the lack of flow. Subsequently, DNAs were stretched by initiating the flow of protein in imaging buffer (50 mL/min). Experiments were conducted using the IX-83 total internal reflection fluorescence (TIRF) microscope (Evident Scientific, Olympus, Waltham, MA) with a 532 nm laser (Coherent, Santa Clara, CA). Recorded images were acquired with the Micro-manager software (87), and DNAs’ regions-of-interest (ROI) were set using FIJI software (88). Custom MATLAB software codes, based on Gaussian fitting, were utilized to determine quantum dot positions. Detailed procedures and the custom MATLAB codes are accessible in our prior publications (85, 86).

### Size exclusion chromatography with multi-angle light scattering (SEC-MALS)

Wild-type ParB and its mutant variants at 0.65 mg ml^-1^ (20 mM) and BSA standard at 2.0 mg ml^-1^ were run at a flow rate of 0.2 mL min^-1^ in running buffer (20 mM Tris at pH 8.0, 350 mM NaCl) on a Cytiva Superdex 200 Increase 200 10/300 GL size exclusion chromatography column. The column was attached to an AKTA Pure fast protein liquid chromatography system (FPLC) coupled with a Wyatt Dawn multi-angle light scattering detector (Wyatt Technology) and a Wyatt Optilab differential refractive index detector (Wyatt Technology). Chromatograms were analyzed using the ASTRA 8 software (Wyatt Technology) to determine the molecular weights and oligomeric states.

### Molecular dynamics simulations

Molecular dynamics (MD) simulations were performed with GROMACS software (version 24.2) (89). The atomistic model of apo CTD was constructed on the available NMR structure (PDB: 5NOC), while the DNA-bound CTD model was constructed using the AlphaFold server (90). The protein was inserted in a cubic box with a buffer width of 1.5 nm. We solvated the CTD with explicit water along with neutralization by chlorine and sodium ions to 0.15 M concentration. To describe proteins, DNA, and ions, we used the AMBER99SB force field (91). The TIP3P model (92) was used to treat the water environment. The energy minimization step was performed with the steepest descent algorithm until the maximum force fell below 1000 kJ/mol/min. Coulomb interactions, Van der Waals interactions, and neighbor list cut-off distance were 1.0 nm. We applied the Verlet cut-off scheme for the neighbor list with the electrostatic interactions calculated by the particle mesh Ewald (93) method with a grid spacing of 0.125 nm. The system was first equilibrated with a position restraint on all protein-heavy atoms to hold a temperature at 308 K with the use of the v-rescale thermostat (94) and then brought to 1 bar pressure with the use of an improved version of the Berendsen barostat (c-rescale) (95). All bonds to hydrogens were constrained with the LINKS algorithm (96). Following the equilibration stage (Extended Data Fig. 4a-f), our 500 ns production run had position restraints removed.

We used truncated average approximation (97) to the Lipari–Szabo model-free approach (44) to calculate the order parameters (S^2^). The S^2^ was determined by taking the average over the second half of each C(t) relaxation curve, where C(t) is the angular autocorrelation function of the second-order Legendre polynomial for the unit vector pointing along the amide bond vector calculated from trajectories generated during the production runs. Average structures were obtained over the time simulation had been conducted.

### Plasmid construction for *in vivo* experiments

The following variants of **pWX563** [*pelB::Psoj-mgfpmut3-spo0J (ΔparS) tet*] (33) were constructed through site-directed mutagenesis as described below.

**pWX1227** [*pelB::Psoj-mgfpmut3-spo0J-A232G(ΔparS) tet*] was constructed by an isothermal assembly reaction containing two fragments: 1) pWX563 amplified using oWX3485 and oWX418; 2) pWX563 amplified with oWX3484 and oWX2071. This procedure introduced the A232G mutation into pWX563. The construct was sequenced by whole plasmid sequencing.

**pWX1228** [*pelB::Psoj-mgfpmut3-spo0J-E261A(ΔparS) tet*] was constructed by an isothermal assembly reaction containing two fragments: 1) pWX563 amplified using oWX3487 and oWX418; 2) pWX563 amplified with oWX3486 and oWX2071. This procedure introduced the E261A mutation into pWX563. The construct was sequenced by whole plasmid sequencing.

**pWX1229** [*pelB::Psoj-mgfpmut3-spo0J-K257A(ΔparS) tet*] was constructed by an isothermal assembly reaction containing two fragments: 1) pWX563 amplified using oWX3489 and oWX418; 2) pWX563 amplified with oWX3488 and oWX2071. This procedure introduced the K257A mutation into pWX563. The construct was sequenced by whole plasmid sequencing.

**pWX1230** [*pelB::Psoj-mgfpmut3-spo0J-K256A(ΔparS) tet*] was constructed by an isothermal assembly reaction containing two fragments: 1) pWX563 amplified using oWX3491 and oWX418; 2) pWX563 amplified with oWX3490 and oWX2071. This procedure introduced the K256A mutation into pWX563. The construct was sequenced by whole plasmid sequencing.

**pWX1323** [*pelB::Psoj-mgfpmut3-spo0J-K252A (ΔparS) tet*] was constructed by an isothermal assembly reaction containing two fragments: 1) pWX563 amplified using oWX3807 and oWX418; 2) pWX563 amplified with oWX3806 and oWX2071. This procedure introduced the K252A mutation into pWX563. The construct was sequenced by whole plasmid sequencing.

**pWX1324** [*pelB::Psoj-mgfpmut3-spo0J-K259A (ΔparS) tet*] was constructed by an isothermal assembly reaction containing two fragments: 1) pWX563 amplified using oWX3809 and oWX418; 2) pWX563 amplified with oWX3808 and oWX2071. This procedure introduced the K259A mutation into pWX563. The construct was sequenced by whole plasmid sequencing.

**pWX1325** [*pelB::Psoj-mgfpmut3-spo0J-K252A-K259A (ΔparS) tet*] was constructed by an isothermal assembly reaction containing two fragments: 1) pWX563 amplified using oWX3811 and oWX418; 2) pWX563 amplified with oWX3810 and oWX2071. This procedure introduced the K252A and K259A mutations into pWX563. The construct was sequenced by whole plasmid sequencing.

**pWX1326** [*pelB::Psoj-mgfpmut3-spo0J-K252A-K255A-K259A (ΔparS) tet*] was constructed by an isothermal assembly reaction containing two fragments: 1) pWX563 amplified using oWX3813 and oWX418; 2) pWX563 amplified with oWX3812 and oWX2071. This procedure introduced the K252A, K255A, and K259A mutations into pWX563. The construct was sequenced by whole plasmid sequencing.

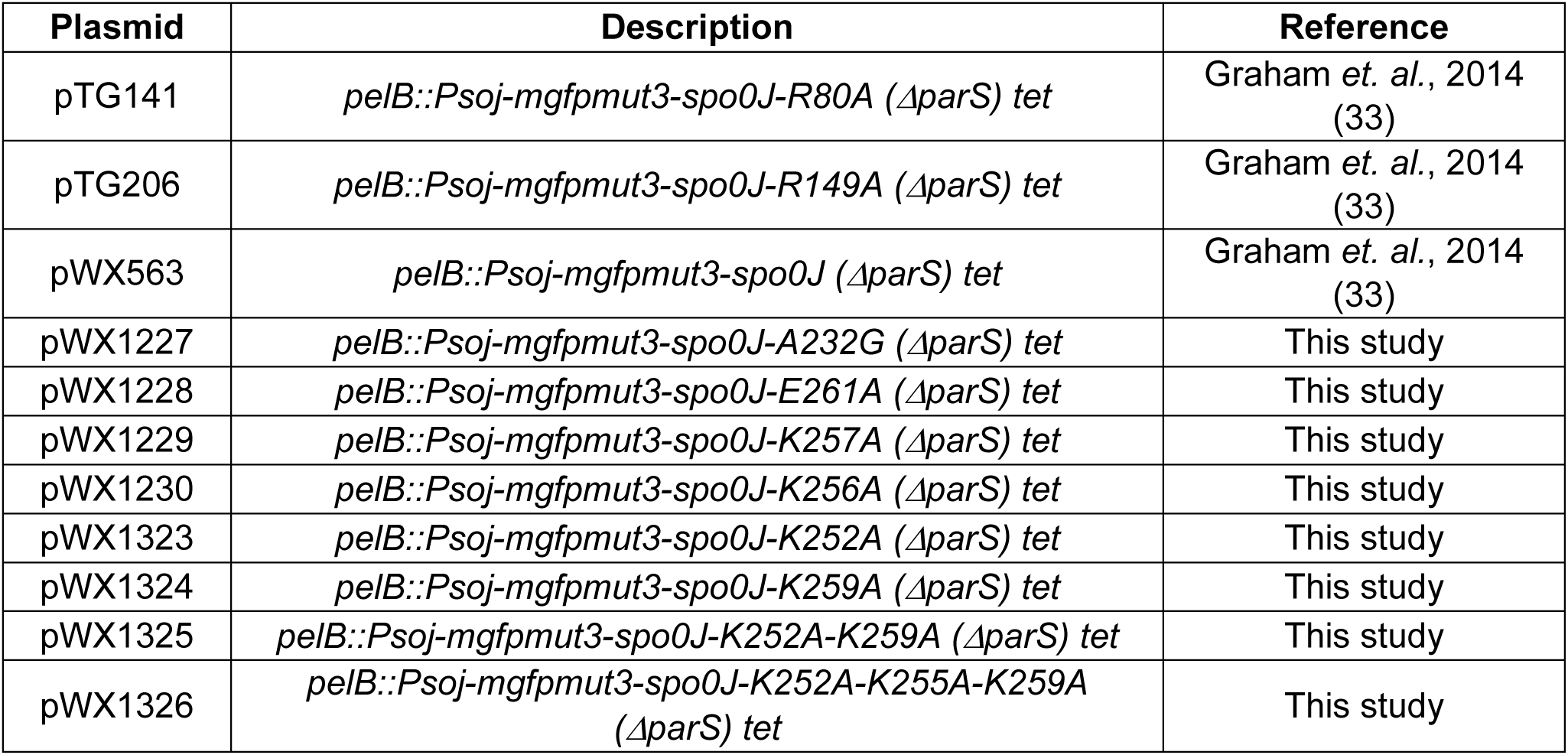

### Strain building

*B. subtilis* strains were derived from the PY79 wild type strain. Strains were built through transformations of plasmids or gDNA. After transformation into *B. subtilis,* gDNA was extracted and the mutated region was amplified using oML85 and oWX507. The amplicon was sequenced by whole plasmid sequencing (linear/amplicon). Strains were grown in defined rich Casein Hydrolysate (CH) medium at 37°C.

| Strain | Genotype | Reference | Figures |
| --- | --- | --- | --- |
| PY79 | wild type | Youngman *et. al.*, 1983 (98) |  |
| BWX2210 (BDR2795) | $\Delta spo0J::spec$, $sacA::hbs-mcherry kan$ | X.W. and D.Z.R., unpublished | |
| BWX2213 (BDR2798) | $\Delta spo0J::spec$, $sacA::hbs-mcherry kan$, $pelB::Psoj-mgfpmut3-spo0J (\Delta parS) tet$ | Graham *et. al.*, 2014 (33) | |
| BWX2214 (BDR2799) | $\Delta spo0J::spec$, $sacA::hbs-mcherry kan$, $pelB::Psoj-mgfpmut3-spo0J-R80A (\Delta parS) tet$ | Graham *et. al.*, 2014 (33) | |
| BWX5754 | $\Delta spo0J::spec$, $sacA::hbs-mcherry kan$, $pelB::Psoj-mgfpmut3-spo0J-A232G (\Delta parS) tet$ | This study | |
| BWX5756 | $\Delta spo0J::spec$, $sacA::hbs-mcherry kan$, $pelB::Psoj-mgfpmut3-spo0J-E261A (\Delta parS) tet$ | This study | |
| BWX5758 | $\Delta spo0J::spec$, $sacA::hbs-mcherry kan$, $pelB::Psoj-mgfpmut3-spo0J-K257A (\Delta parS) tet$ | This study | |
| BWX5760 | $\Delta spo0J::spec$, $sacA::hbs-mcherry kan$, $pelB::Psoj-mgfpmut3-spo0J-K256A (\Delta parS) tet$ | This study | |
| BWX6087 | $\Delta spo0J::spec$, $sacA::hbs-mcherry kan$, $pelB::Psoj-mgfpmut3-spo0J-K252A (\Delta parS) tet$ | This study | |
| BWX6089 | $\Delta spo0J::spec$, $sacA::hbs-mcherry kan$, $pelB::Psoj-mgfpmut3-spo0J-K259A (\Delta parS) tet$ | This study | |
| BWX6091 | $\Delta spo0J::spec$, $sacA::hbs-mcherry kan$, $pelB::Psoj-mgfpmut3-spo0J-K252A-K259A (\Delta parS) tet$ | This study | |
| BWX6093 | $\Delta spo0J::spec$, $sacA::hbs-mcherry kan$, $pelB::Psoj-mgfpmut3-spo0J-K252A-K255A-K259A (\Delta parS) tet$ | This study | |

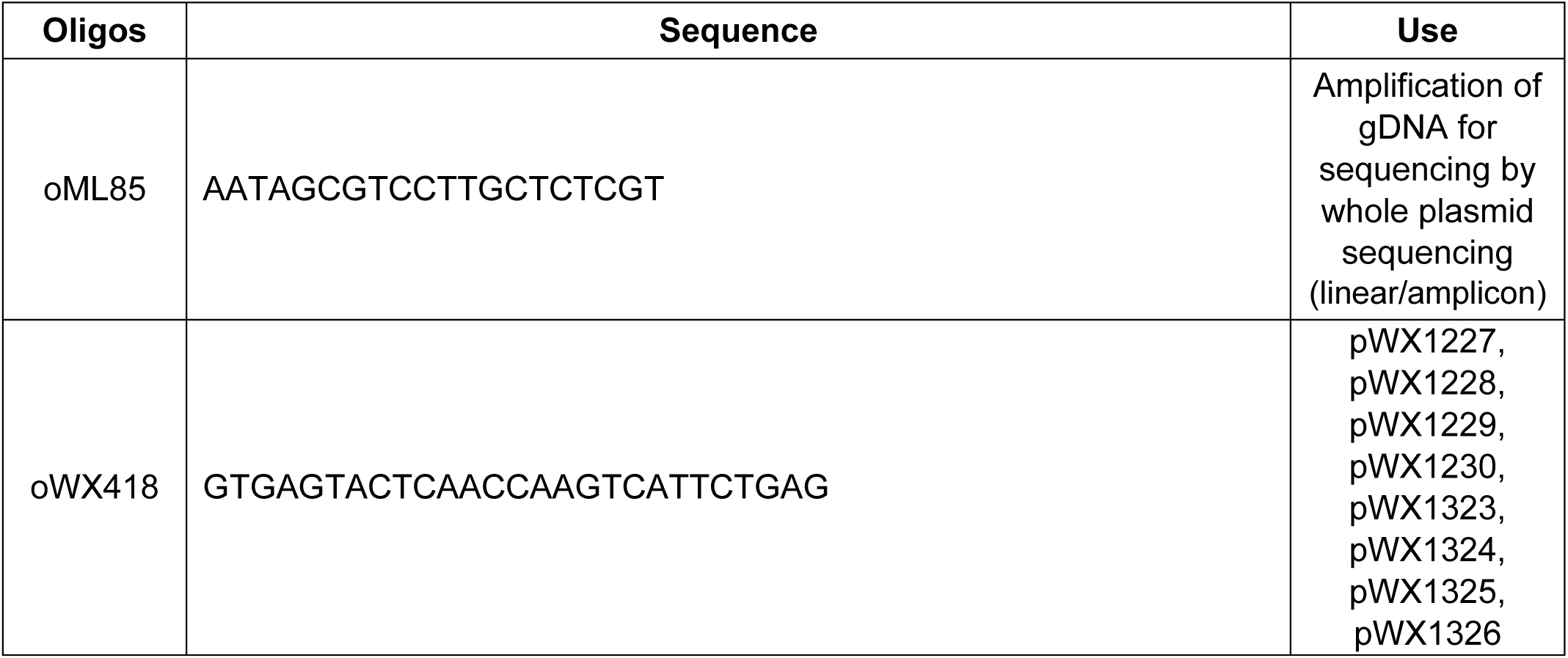
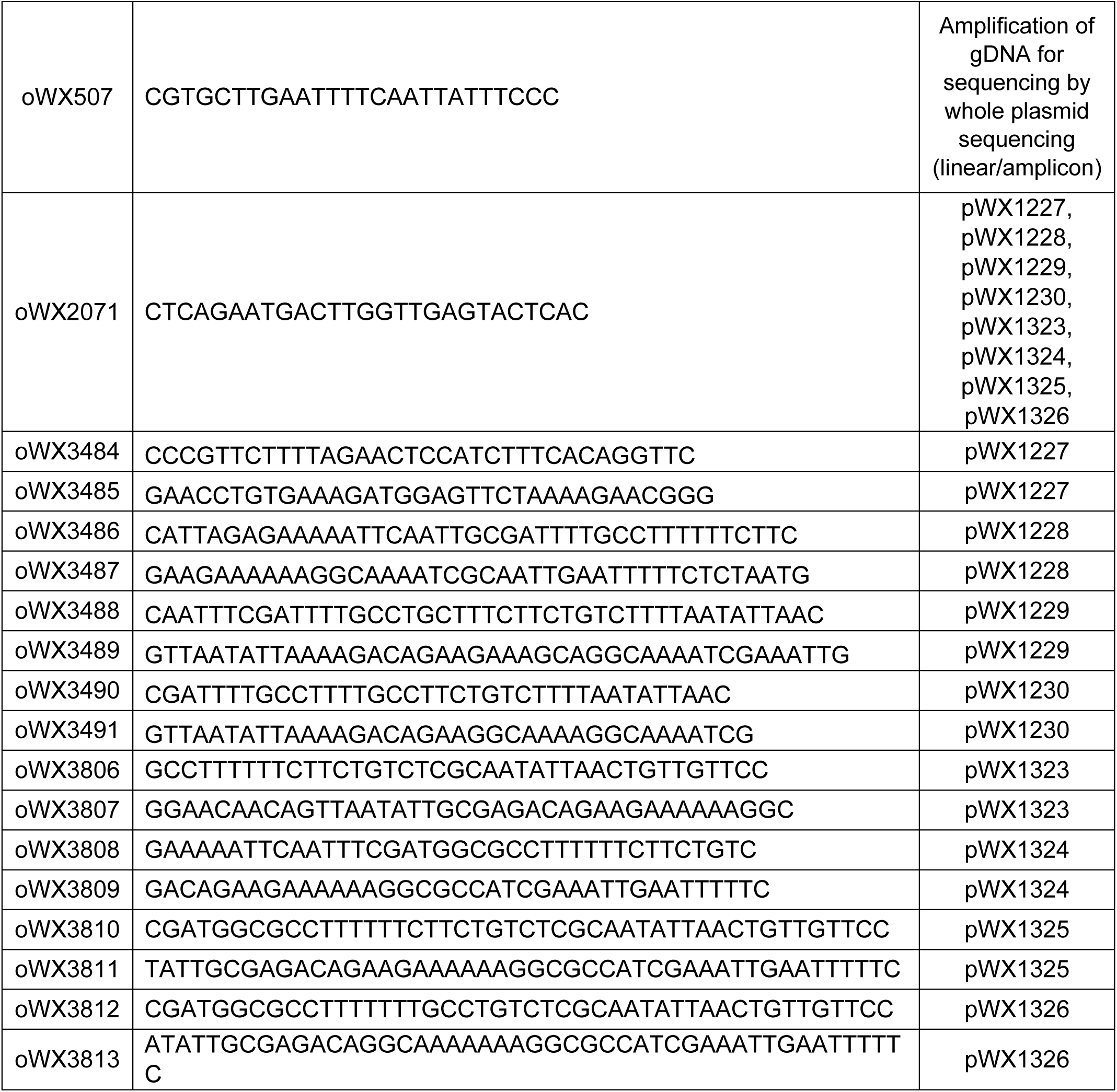

### Fluorescence microscopy

Strains were imaged during exponential phase using a Nikon Ti2 microscope with a Plan Apo 100x/1.45NA phase contrast oil objective and an sCMOS camera. Images were cropped and adjusted using NIS-Elements Advanced Research Imaging Software. Figures were created in Adobe Illustrator.

### Heat resistance sporulation assay

Sporulation efficiency was determined as described by Harwood and Cutting (99). Briefly, a freshly streaked colony was inoculated in 2 mL of Difco Sporulation Medium (DSM) and incubated for 24 hours in a roller drum at 37°C. The cultures were serially diluted by 10-fold in the dilution medium containing T-Base and 1 mM MgSO4. To determine total colony forming units, 200 μL of the 10^-6^ dilution and 50 μL of the 10^-5^ dilution were plated on separate DSM agar plates. To kill the non-sporulated cells, the diluted cultures were incubated at 80°C for 20 minutes. 100 µl of the heat-killed dilutions were plated onto DSM agar plates. All plates were incubated overnight at 37°C.

## Notes

### Competing Interest Statement

The authors have declared no competing interest.

## REFERENCES

1. Leonard, T.A., Butler, P.J. and Löwe, J. (2005) Bacterial chromosome segregation: structure and DNA binding of the Soj dimer--a conserved biological switch. EMBO J, 24, 270–282.

2. Roberts, M.A.J., Wadhams, G.H., Hadfield, K.A., Tickner, S. and Armitage, J.P. (2012) ParA-like protein uses nonspecific chromosomal DNA binding to partition protein complexes. Proc Natl Acad Sci U S A, 109, 6698–6703.

3. Lin, D.C.H. and Grossman, A.D. (1998) Identification and characterization of a bacterial chromosome partitioning site. Cell, 92, 675–685.

4. Breier, A.M. and Grossman, A.D. (2007) Whole-genome analysis of the chromosome partitioning and sporulation protein Spo0J (ParB) reveals spreading and origin-distal sites on the Bacillus subtilis chromosome. Mol Microbiol, 64, 703–718.

5. Livny, J., Yamaichi, Y. and Waldor, M.K. (2007) Distribution of centromere-like parS sites in bacteria: insights from comparative genomics. J Bacteriol, 189, 8693–8703.

6. Ptacin, J.L., Lee, S.F., Garner, E.C., Toro, E., Eckart, M., Comolli, L.R., Moerner, W.E. and Shapiro, L. (2010) A spindle-like apparatus guides bacterial chromosome segregation. Nat Cell Biol, 12, 791–798.

7. Lim, H.C., Surovtsev, I.V., Beltran, B.G., Huang, F., Bewersdorf, J. and Jacobs-Wagner, C. (2014) Evidence for a DNA-relay mechanism in ParABS-mediated chromosome segregation. Elife, 3, e02758.

8. Surovtsev, I. V, Campos, M. and Jacobs-Wagner, C. (2016) DNA-relay mechanism is sufficient to explain ParA-dependent intracellular transport and patterning of single and multiple cargos. Proc Natl Acad Sci U S A, 113, E7268–E7276.

9. Hu, L., Vecchiarelli, A.G., Mizuuchi, K., Neuman, K.C. and Liu, J. (2017) Brownian Ratchet Mechanism for Faithful Segregation of Low-Copy-Number Plasmids. Biophys J, 112, 1489–1502.

10. Sullivan, N.L., Marquis, K.A. and Rudner, D.Z. (2009) Recruitment of SMC by ParB-parS Organizes the Origin Region and Promotes Efficient Chromosome Segregation. Cell, 137, 697–707.

11. Gruber, S. and Errington, J. (2009) Recruitment of Condensin to Replication Origin Regions by ParB/SpoOJ Promotes Chromosome Segregation in B. subtilis. Cell, 137, 685–696.

12. Murray, H. and Errington, J. (2008) Dynamic control of the DNA replication initiation protein DnaA by Soj/ParA. Cell, 135, 74–84.

13. Jalal, A.S.B. and Le, T.B.K. (2020) Bacterial chromosome segregation by the ParABS system. Open Biol, 10, 200097.

14. Tišma, M., Kaljević, J., Gruber, S., Le, T.B.K. and Dekker, C. (2024) Connecting the dots: key insights on ParB for chromosome segregation from single-molecule studies. FEMS Microbiol Rev, 48, 1–13.

15. Mohl, D.A. and Gober, J.W. (1997) Cell cycle-dependent polar localization of chromosome partitioning proteins in Caulobacter crescentus. Cell, 88, 675–684.

16. Ireton, K., Gunther IV, N.W. and Grossman, A.D. (1994) spo0J is required for normal chromosome segregation as well as the initiation of sporulation in Bacillus subtilis. J Bacteriol, 176, 5320–5329.

17. Soh, Y.M., Bürmann, F., Shin, H.C., Oda, T., Jin, K.S., Toseland, C.P., Kim, C., Lee, H., Kim, S.J., Kong, M.S., et al. (2015) Molecular basis for SMC rod formation and its dissolution upon DNA binding. Mol Cell, 57, 290– 303.

18. Osorio-Valeriano, M., Altegoer, F., Steinchen, W., Urban, S., Liu, Y., Bange, G. and Thanbichler, M. (2019) ParB-type DNA Segregation Proteins Are CTP-Dependent Molecular Switches. Cell, 179, 1512–1524.e15.

19. Osorio-Valeriano, M., Altegoer, F., Das, C.K., Steinchen, W., Panis, G., Connolley, L., Giacomelli, G., Feddersen, H., Corrales-Guerrero, L., Giammarinaro, P.I., et al. (2021) The CTPase activity of ParB determines the size and dynamics of prokaryotic DNA partition complexes. Mol Cell, 81, 3992–4007.

20. Jalal, A.S., Tran, N.T., Stevenson, C.E., Chimthanawala, A., Badrinarayanan, A., Lawson, D.M. and Le, T.B. (2021) A CTP-dependent gating mechanism enables ParB spreading on DNA. Elife, 10, e69676.

21. Davis, M.A., Martin, K.A. and Austin, S.J. (1992) Biochemical activities of the parA partition protein of the P1 plasmid. Mol Microbiol, 6, 1141–1147.

22. Radnedge, L., Youngren, B., Davis, M. and Austin, S. (1998) Probing the structure of complex macromolecular interactions by homolog specificity scanning: the P1 and P7 plasmid partition systems. EMBO J, 17, 6076–6085.

23. Zhao, Y., Guo, L., Hu, J., Ren, Z., Li, Y., Hu, M., Zhang, X., Bi, L., Li, D., Ma, H., et al. (2024) Phase-separated ParB enforces diverse DNA compaction modes and stabilizes the parS-centered partition complex. Nucleic Acids Res, 52, 8385–8398.

24. Leonard, T.A., Butler, P.J.G. and Löwe, J. (2004) Structural analysis of the chromosome segregation protein Spo0J from Thermus thermophilus. Mol Microbiol, 53, 419–432.

25. Schumacher, M.A. and Funnell, B.E. (2005) Structures of ParB bound to DNA reveal mechanism of partition complex formation. Nature, 438, 516–519.

26. Soh, Y.M., Davidson, I.F., Zamuner, S., Basquin, J., Bock, F.P., Taschner, M., Veening, J.W., de Los Rios, P., Peters, J.M. and Gruber, S. (2019) Self-organization of parS centromeres by the ParB CTP hydrolase. Science, 366, 1129–1133.

27. Chen, B.-W., Lin, M.-H., Chu, C.-H., Hsu, C.-E. and Sun, Y.-J. (2015) Insights into ParB spreading from the complex structure of Spo0J and parS. Proc Natl Acad Sci U S A, 112, 6613–6618.

28. Taylor, J.A., Pastrana, C.L., Butterer, A., Pernstich, C., Gwynn, E.J., Sobott, F., Moreno-Herrero, F. and Dillingham, M.S. (2015) Specific and non-specific interactions of ParB with DNA: Implications for chromosome segregation. Nucleic Acids Res, 43, 719–731.

29. Fisher, G.L.M., Pastrana, C.L., Higman, V.A., Koh, A., Taylor, J.A., Butterer, A., Craggs, T., Sobott, F., Murray, H., Crump, M.P., et al. (2017) The structural basis for dynamic DNA binding and bridging interactions which condense the bacterial centromere. Elife, 6, e28086.

30. Murray, H., Ferreira, H. and Errington, J. (2006) The bacterial chromosome segregation protein Spo0J spreads along DNA from parS nucleation sites. Mol Microbiol, 61, 1352–1361.

31. Tran, N.T., Stevenson, C.E., Som, N.F., Thanapipatsiri, A., Jalal, A.S.B. and Le, T.B.K. (2018) Permissive zones for the centromere-binding protein ParB on the Caulobacter crescentus chromosome. Nucleic Acids Res, 46, 1196–1209.

32. Rodionov, O., ŁObocka, M. and Yarmolinsky, M. (1999) Silencing of genes flanking the P1 plasmid centromere. Science, 283, 546–549.

33. Graham, T.G.W., Wang, X., Song, D., Etson, C.M., van Oijen, A.M., Rudner, D.Z. and Loparo, J.J. (2014) ParB spreading requires DNA bridging. Genes Dev, 28, 1228–1238.

34. Sanchez, A., Cattoni, D.I., Walter, J.-C., Rech, J., Parmeggiani, A., Nollmann, M. and Bouet, J.-Y. (2015) Stochastic Self-Assembly of ParB Proteins Builds the Bacterial DNA Segregation Apparatus. Cell Syst, 1, 163–173.

35. Antar, H., Soh, Y.-M., Zamuner, S., Bock, F.P., Anchimiuk, A., De Los Rios, P. and Gruber, S. (2021) Relief of ParB autoinhibition by parS DNA catalysis and recycling of ParB by CTP hydrolysis promote bacterial centromere assembly. Sci Adv, 7, esbj2854.

36. Jalal, A.S., Tran, N.T. and Le, T.B. (2020) ParB spreading on DNA requires cytidine triphosphate in vitro. Elife, 9, 1–24.

37. Balaguer, F. de A., Aicart-Ramos, C., Fisher, G.L., de Bragança, S., Martin-Cuevas, E.M., Pastrana, C.L., Dillingham, M.S. and Moreno-Herrero, F. (2021) CTP promotes efficient ParB-dependent DNA condensation by facilitating one-dimensional diffusion from parS. Elife, 10, e677554.

38. Guo, L., Zhao, Y., Zhang, Q., Feng, Y., Bi, L., Zhang, X., Wang, T., Liu, C., Ma, H. and Sun, B. (2022) Stochastically multimerized ParB orchestrates DNA assembly as unveiled by single-molecule analysis. Nucleic Acids Res, 50, 9294–9305.

39. Jarymowycz, V.A. and Stone, M.J. (2006) Fast time scale dynamics of protein backbones: NMR relaxation methods, applications, and functional consequences. Chem Rev, 106, 1624–1671.

40. Kay, L.E., Torchia, D.A. and Bax, A. (1989) Backbone dynamics of proteins as studied by nitrogen-15 inverse detected heteronuclear NMR spectroscopy: application to staphylococcal nuclease. Biochemistry, 28, 8972–8979.

41. Clore, G.M., Driscoll, P.C., Wingfield, P.T. and Gronenborn, A.M. (1990) Analysis of the backbone dynamics of interleukin-1 beta using two-dimensional inverse detected heteronuclear 15N-1H NMR spectroscopy. Biochemistry, 29, 7387–7401.

42. Tjandra, N., Feller, S.E., Pastor, R.W. and Bax, A. (1995) Rotational diffusion anisotropy of human ubiquitin from 15N NMR relaxation. J Am Chem Soc, 117, 12562–12566.

43. Tjandra, N., Garrett, D.S., Gronenborn, A.M., Bax, A. and Clore, G.M. (1997) Defining long range order in NMR structure determination from the dependence of heteronuclear relaxation times on rotational diffusion anisotropy. Nat Struct Biol, 4, 443–449.

44. Lipari, G. and Szabo, A. (1982) Model-free approach to the interpretation of nuclear magnetic resonance relaxation in macromolecules. 1. Theory and range of validity. J Am Chem Soc, 104, 4546–4559.

45. Sapienza, P.J. and Lee, A.L. (2010) Using NMR to study fast dynamics in proteins: methods and applications. Curr Opin Pharmacol, 10, 723–730.

46. Song, D., Rodrigues, K., Graham, T.G.W. and Loparo, J.J. (2017) A network of cis and trans interactions is required for ParB spreading. Nucleic Acids Res, 45, 7106–7117.

47. Case, D.A. (2002) Molecular Dynamics and NMR Spin Relaxation in Proteins. Acc Chem Res, 35, 325–331.

48. Hornak, V., Abel, R., Okur, A., Strockbine, B., Roitberg, A. and Simmerling, C. (2006) Comparison of multiple Amber force fields and development of improved protein backbone parameters. Proteins, 65, 712–725.

49. Williamson, M.P. (2013) Using chemical shift perturbation to characterise ligand binding. Prog Nucl Magn Reson Spectrosc, 73, 1–16.

50. Shehzad, S. and Kim, H. (2025) Single-molecule DNA-flow stretching assay as a versatile hybrid tool for investigating DNA-protein interactions. BMB Rep, 58, 41–51.

51. Molina, M., Way, L.E., Ren, Z., Liao, Q., Guerra, B., Shields, B., Wang, X. and Kim, H. (2023) A framework to validate fluorescently labeled DNA-binding proteins for single-molecule experiments. Cell Reports Methods, 3, 100614.

52. Tišma, M., Panoukidou, M., Antar, H., Soh, Y.-M., Barth, R., Pradhan, B., Barth, A., Van Der Torre, J., Michieletto, D., Gruber, S., et al. (2022) ParB proteins can bypass DNA-bound roadblocks via dimer-dimer recruitment. Sci Adv, 8, esbn3299.

53. Broedersz, C.P., Wang, X., Meir, Y., Loparo, J.J., Rudner, D.Z. and Wingreen, N.S. (2014) Condensation and localization of the partitioning protein ParB on the bacterial chromosome. Proc Natl Acad Sci U S A, 111, 8809–8814.

54. Walter, J.-C., Walliser, N.-O., David, G., Dorignac, J., Geniet, F., Palmeri, J., Parmeggiani, A., Wingreen, N.S. and Broedersz, C.P. (2018) Looping and clustering model for the organization of protein-DNA complexes on the bacterial genome. New J Phys, 20, 035002.

55. Sharpe, M.E. and Errington, J. (1996) The Bacillus subtilis soj-spo0J locus is required for a centromere-like function involved in prespore chromosome partitioning. Mol Microbiol, 21, 501–509.

56. Lin, D.C.H., Levin, P.A. and Grossman, A.D. (1997) Bipolar localization of a chromosome partition protein in Bacillus subtilis. Proc Natl Acad Sci U S A, 94, 4721–4726.

57. Webb, C.D., Teleman, A., Gordon, S., Straight, A., Belmont, A., Lin, D.C.H., Grossman, A.D., Wright, A. and Losick, R. (1997) Bipolar localization of the replication origin regions of chromosomes in vegetative and sporulating cells of B. subtilis. Cell, 88, 667–674.

58. Autret, S., Nair, R. and Errington, J. (2001) Genetic analysis of the chromosome segregation protein Spo0J of Bacillus subtilis: evidence for separate domains involved in DNA binding and interactions with Soj protein. Mol Microbiol, 41, 743–55.

59. Henzler-Wildman, K. and Kern, D. (2007) Dynamic personalities of proteins. Nature, 450, 964–972.

60. Kalodimos, C.G., Biris, N., Bonvin, A.M.J.J., Levandoski, M.M., Guennuegues, M., Boelens, R. and Kaptein, R. (2004) Structure and flexibility adaptation in nonspecific and specific protein-DNA complexes. Science, 305, 386–389.

61. Sullivan, S.M. and Maddock, J.R. (2000) Bacterial sporulation: pole-to-pole protein oscillation. Current biology, 10, R159–R161.

62. Marston, A.L. and Errington, J. (1999) Dynamic movement of the ParA-like Soj protein of B. subtilis and its dual role in nucleoid organization and developmental regulation. Mol Cell, 4, 673–682.

63. Walter, J.-C., Rech, J., Walliser, N.-O., Dorignac, J., Geniet, F., Palmeri, J., Parmeggiani, A. and Bouet, J.-Y. (2020) Physical Modeling of a Sliding Clamp Mechanism for the Spreading of ParB at Short Genomic Distance from Bacterial Centromere Sites. iScience, 23, 101861.

64. Moldovan, G.-L., Pfander, B. and Jentsch, S. (2007) PCNA, the maestro of the replication fork. Cell, 129, 665–679.

65. Krishna, T.S., Kong, X.P., Gary, S., Burgers, P.M. and Kuriyan, J. (1994) Crystal structure of the eukaryotic DNA polymerase processivity factor PCNA. Cell, 79, 1233–1243.

66. Gulbis, J.M., Kelman, Z., Hurwitz, J., O’Donnell, M. and Kuriyan, J. (1996) Structure of the C-terminal region of p21(WAF1/CIP1) complexed with human PCNA. Cell, 87, 297–306.

67. De March, M., Merino, N., Barrera-Vilarmau, S., Crehuet, R., Onesti, S., Blanco, F.J. and De Biasio, A. (2017) Structural basis of human PCNA sliding on DNA. Nat Commun, 8, 13935.

68. Wing, R., Drew, H., Takano, T., Broka, C., Tanaka, S., Itakura, K. and Dickerson, R.E. (1980) Crystal structure analysis of a complete turn of B-DNA. Nature, 287, 755–758.

69. Fukuda, K., Morioka, H., Imajou, S., Ikeda, S., Ohtsuka, E. and Tsurimoto, T. (1995) Structure-function relationship of the eukaryotic DNA replication factor, proliferating cell nuclear antigen. J Biol Chem, 270, 22527–22534.

70. Zhou, Y. and Hingorani, M.M. (2012) Impact of individual proliferating cell nuclear antigen-DNA contacts on clamp loading and function on DNA. J Biol Chem, 287, 35370–35381.

71. Abramson, J., Adler, J., Dunger, J., Evans, R., Green, T., Pritzel, A., Ronneberger, O., Willmore, L., Ballard, A.J., Bambrick, J., et al. (2024) Accurate structure prediction of biomolecular interactions with AlphaFold 3. Nature, 630, 493–500.

72. Lee, R., Yang, K. and Lee, J.-B. (2019) Multiplexed single-molecule flow-stretching bead assay for DNA enzymology. BMB Rep, 52, 589–594.

73. Cavanagh, J., Fairbrother, W.J., Palmer, A.G.I., Rance, M. and Skelton, N.J. (2007) Protein NMR Spectroscopy Elsevier.

74. Vold, R.L., Waugh, J.S., Klein, M.P. and Phelps, D.E. (1968) Measurement of Spin Relaxation in Complex Systems. J Chem Phys, 48, 3831–3832.

75. Meiboom, S. and Gill, D. (1958) Modified Spin-Echo Method for Measuring Nuclear Relaxation Times. Review of Scientific Instruments, 29, 688–691.

76. Noggle, J.H. and Schirmer, R.E. (1971) The Nuclear Overhauser Effect: Chemical Applications Academic Press, New York.

77. Kay, L.E., Nicholson, L.K., Delaglio, F., Bax, A. and Torchia, D.A. (1992) Pulse sequences for removal of the effects of cross correlation between dipolar and chemical-shift anisotropy relaxation mechanisms on the measurement of heteronuclear T1 and T2 values in proteins. Journal of Magnetic Resonance (1969), 97, 359–375.

78. Barbato, G., Ikura, M., Kay, L.E., Pastor, R.W. and Bax, A. (1992) Backbone dynamics of calmodulin studied by nitrogen-15 relaxation using inverse detected two-dimensional NMR spectroscopy: the central helix is flexible. Biochemistry, 31, 5269–5278.

79. Palmer, A.G., Skelton, N.J., Chazin, W.J., Wright, P.E. and Rance, M. (1992) Suppression of the effects of cross-correlation between dipolar and anisotropic chemical shift relaxation mechanisms in the measurement of spin-spin relaxation rates. Mol Phys, 75, 699–711.

80. Delaglio, F., Grzesiek, S., Vuister, Geerten W., Zhu, G., Pfeifer, J. and Bax, A. (1995) NMRPipe: A multidimensional spectral processing system based on UNIX pipes. J Biomol NMR, 6, 277–293.

81. Lee, L.K., Rance, M., Chazin, W.J. and Palmer III, A.G. (1997) Rotational diffusion anisotropy of proteins from simultaneous analysis of 15N and 13Cα nuclear spin relaxation. J Biomol NMR, 9, 287–298.

82. Cole, R. and Loria, J.P. (2003) FAST-Modelfree: a program for rapid automated analysis of solution NMR spin-relaxation data. J Biomol NMR, 26, 203–213.

83. Mandel, A.M., Akke, M. and Palmer III, A.G. (1995) Backbone dynamics of Escherichia coli ribonuclease HI: correlations with structure and function in an active enzyme. J Mol Biol, 246, 144–163.

84. Palmer III, A.G., Cavanagh, J., Wright, P.E. and Rance, M. (1991) Sensitivity improvement in proton-detected two-dimensional heteronuclear correlation NMR spectroscopy. Journal of Magnetic Resonance (1969), 93, 151–170.

85. Kim, H. and Loparo, J.J. (2018) Observing Bacterial Chromatin Protein-DNA Interactions by Combining DNA Flow-Stretching with Single-Molecule Imaging. Methods in Molecular Biology, 1837, 277–299.

86. Kim, H. and Loparo, J.J. (2016) Multistep assembly of DNA condensation clusters by SMC. Nat Commun, 7, 1–12.

87. Edelstein, A.D., Tsuchida, M.A., Amodaj, N., Pinkard, H., Vale, R.D. and Stuurman, N. (2014) Advanced methods of microscope control using μManager software. J Biol Methods, 1, e10.

88. Schindelin, J., Arganda-Carreras, I., Frise, E., Kaynig, V., Longair, M., Pietzsch, T., Preibisch, S., Rueden, C., Saalfeld, S., Schmid, B., et al. (2012) Fiji: An open-source platform for biological-image analysis. Nat Methods, 9, 676–682.

89. Abraham, M.J., Murtola, T., Schulz, R., Páll, S., Smith, J.C., Hess, B. and Lindahl, E. (2015) GROMACS: High performance molecular simulations through multi-level parallelism from laptops to supercomputers. SoftwareX, **1–2**, 19–25.

90. Abramson, J., Adler, J., Dunger, J., Evans, R., Green, T., Pritzel, A., Ronneberger, O., Willmore, L., Ballard, A.J., Bambrick, J., et al. (2024) Accurate structure prediction of biomolecular interactions with AlphaFold 3. Nature, 630, 493–500.

91. Hornak, V., Abel, R., Okur, A., Strockbine, B., Roitberg, A. and Simmerling, C. (2006) Comparison of multiple Amber force fields and development of improved protein backbone parameters. *Proteins: Structure*, Function, and Bioinformatics, 65, 712–725.

92. Jorgensen, W.L., Chandrasekhar, J., Madura, J.D., Impey, R.W. and Klein, M.L. (1983) Comparison of simple potential functions for simulating liquid water. J Chem Phys, 79, 926–935.

93. Essmann, U., Perera, L., Berkowitz, M.L., Darden, T., Lee, H. and Pedersen, L.G. (1995) A smooth particle mesh Ewald method. J Chem Phys, 103, 8577–8593.

94. Bussi, G., Donadio, D. and Parrinello, M. (2007) Canonical sampling through velocity rescaling. J Chem Phys, 126, 014101.

95. Bernetti, M. and Bussi, G. (2020) Pressure control using stochastic cell rescaling. J Chem Phys, 153, 114107.

96. Hess, B., Bekker, H., Berendsen, H.J.C. and Fraaije, J.G.E.M. (1997) LINCS: A linear constraint solver for molecular simulations. J Comput Chem, 18, 1463–1472.

97. Bowman, G.R. (2016) Accurately modeling nanosecond protein dynamics requires at least microseconds of simulation. J Comput Chem, 37, 558–566.

98. Youngman, P.J., Perkins, J.B. and Losick, R. (1983) Genetic transposition and insertional mutagenesis in Bacillus subtilis with Streptococcus faecalis transposon Tn917.

99. Harwood, C.R. and Cutting, S.M. (1990) Molecular Biological Methods for Bacillus Wiley.

