## Extended Figures for "ParB C-terminal lysine residues are essential for dimerization, *in vitro* DNA sliding and *in vivo* function"

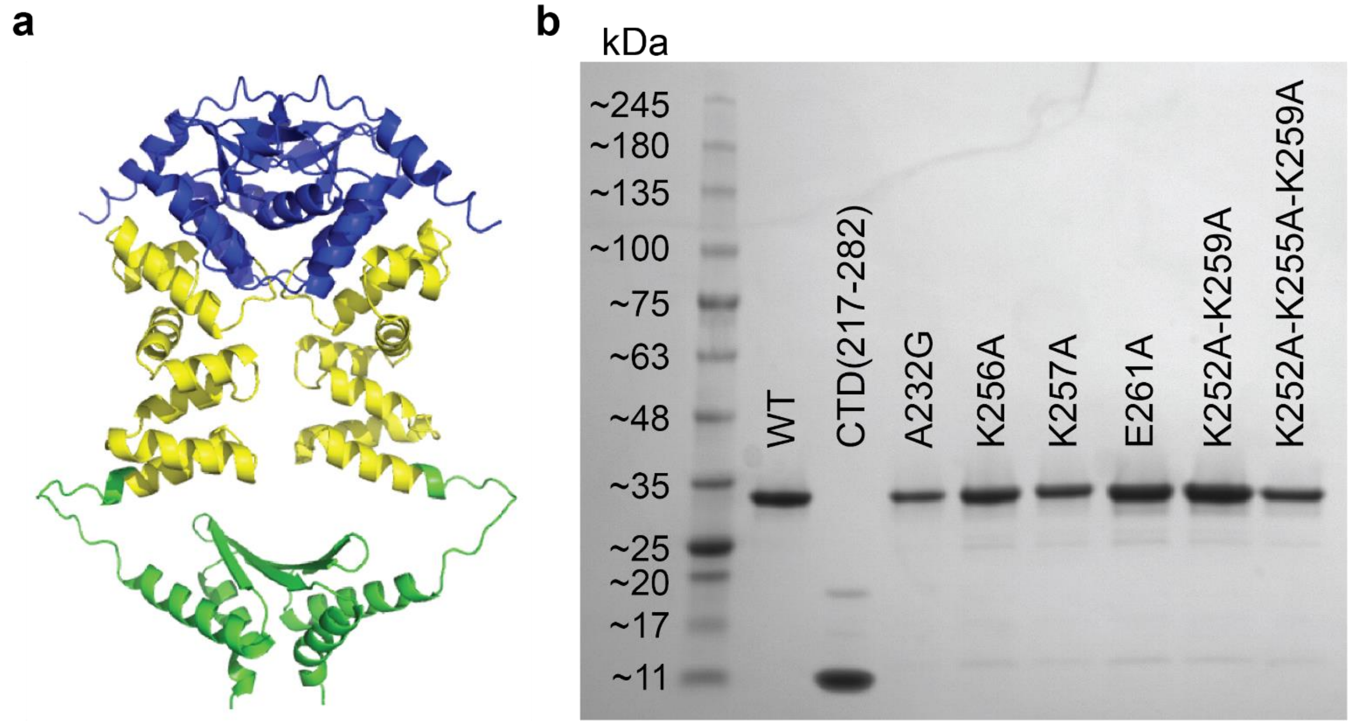


**Extended Data Fig. 1 | *Bacillus subtilis* ParB used in this study.**

**a,** AlphaFold 2 prediction of *B. subtilis* ParB structure. Three distinct domains are shown in colors. Blue: N-terminal domain (NTD), Yellow: Middle DNA-binding domain (DBD), Green: C-terminal domain (CTD)

**b,** SDS-PAGE gel image for a protein ladder (left) and purified *Bacillus subtilis* ParB (BsParB) proteins used in this study.

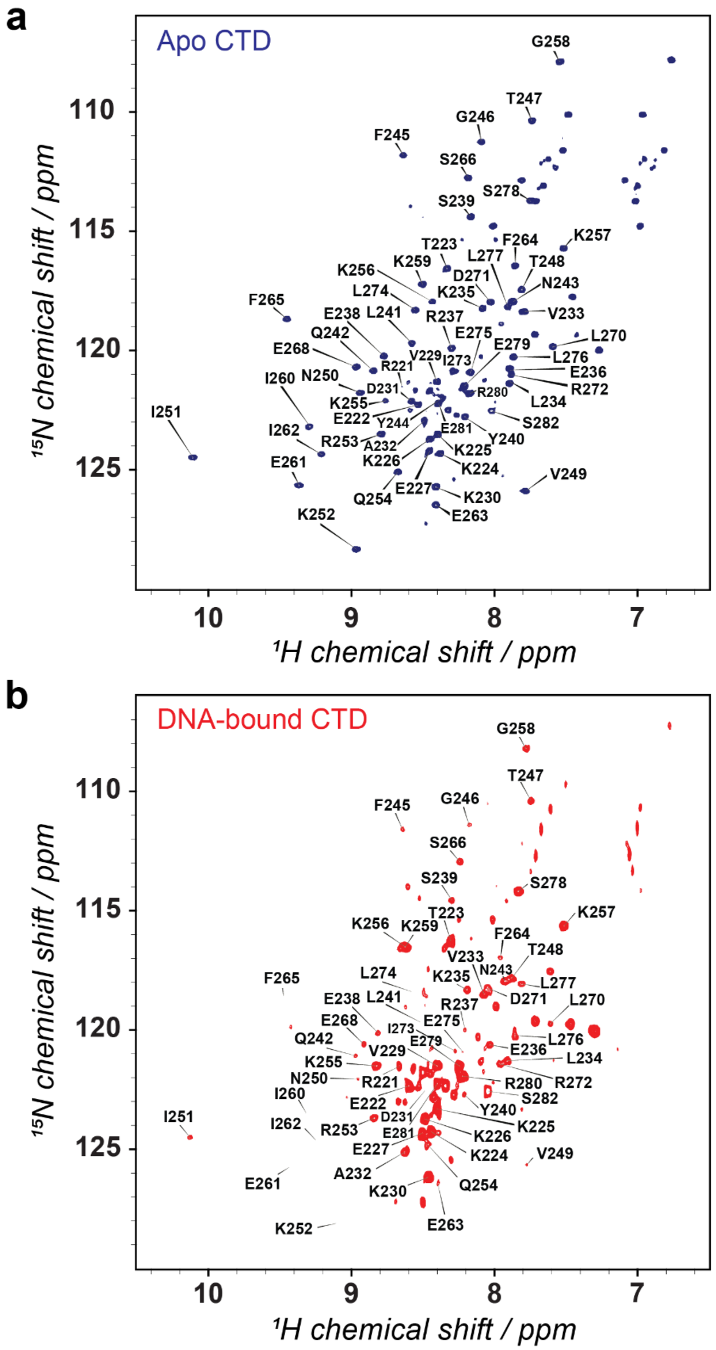


**Extended Data Fig. 2 | $^{1}$H-$^{15}$N heteronuclear single quantum coherence (HSQC) spectra of the *Bacillus subtilis* ParB C-terminal domain (CTD).**

**a-b,** $^{1}$H-$^{15}$N HSQC spectra of apo-CTD and 1.25 equivalent DNA-bound CTD ([DNA] / [CTD] = 1.25), respectively, at 11.7 T. 400 μM CTD in PBS buffer, pH 6.1 at 308 K. While most signals were resolved, the significant exchange broadening led to non-observable signals from G216, Q217, and N218.

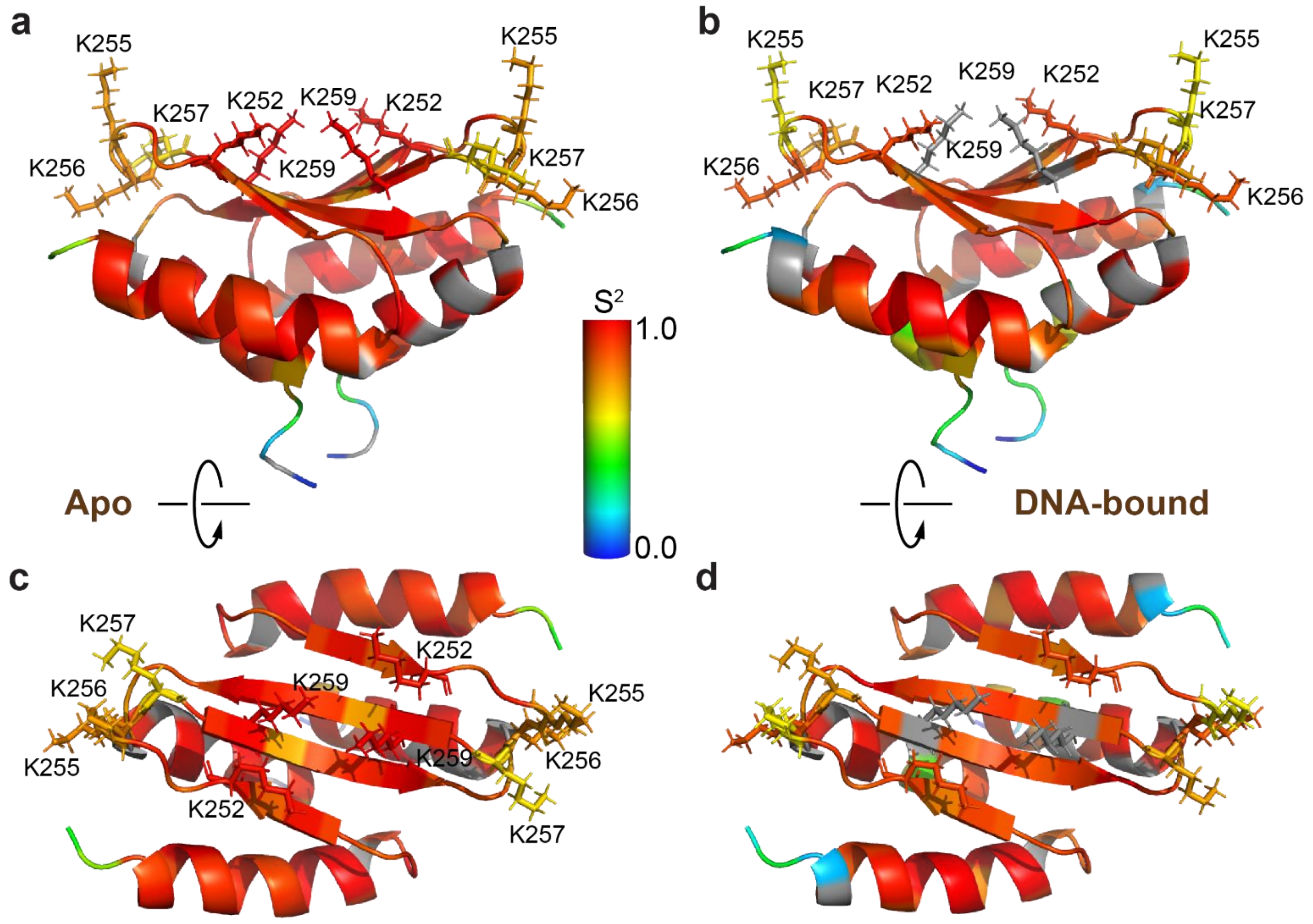


**Extended Data Fig. 3 | Experimentally determined $S^2$ mapped onto an NMR structure of CTD (PDB entry 5NOC).**

**a,b,** Color-coded $S^2$ values from NMR relaxation experiments for (**a**) apo and (**b**) DNA-bound CTD, respectively. Lysines of interest are labeled. These two panels are identical to Fig. 3**a, b** panels.

**c-d,** Views from the top by rotating (**a**) and (**b**) by 90°, respectively. Color scheme: gradient from blue to red for residues with $S^2$ between 0.0 and 1.0, and gray for residues with no assigned $S^2$.

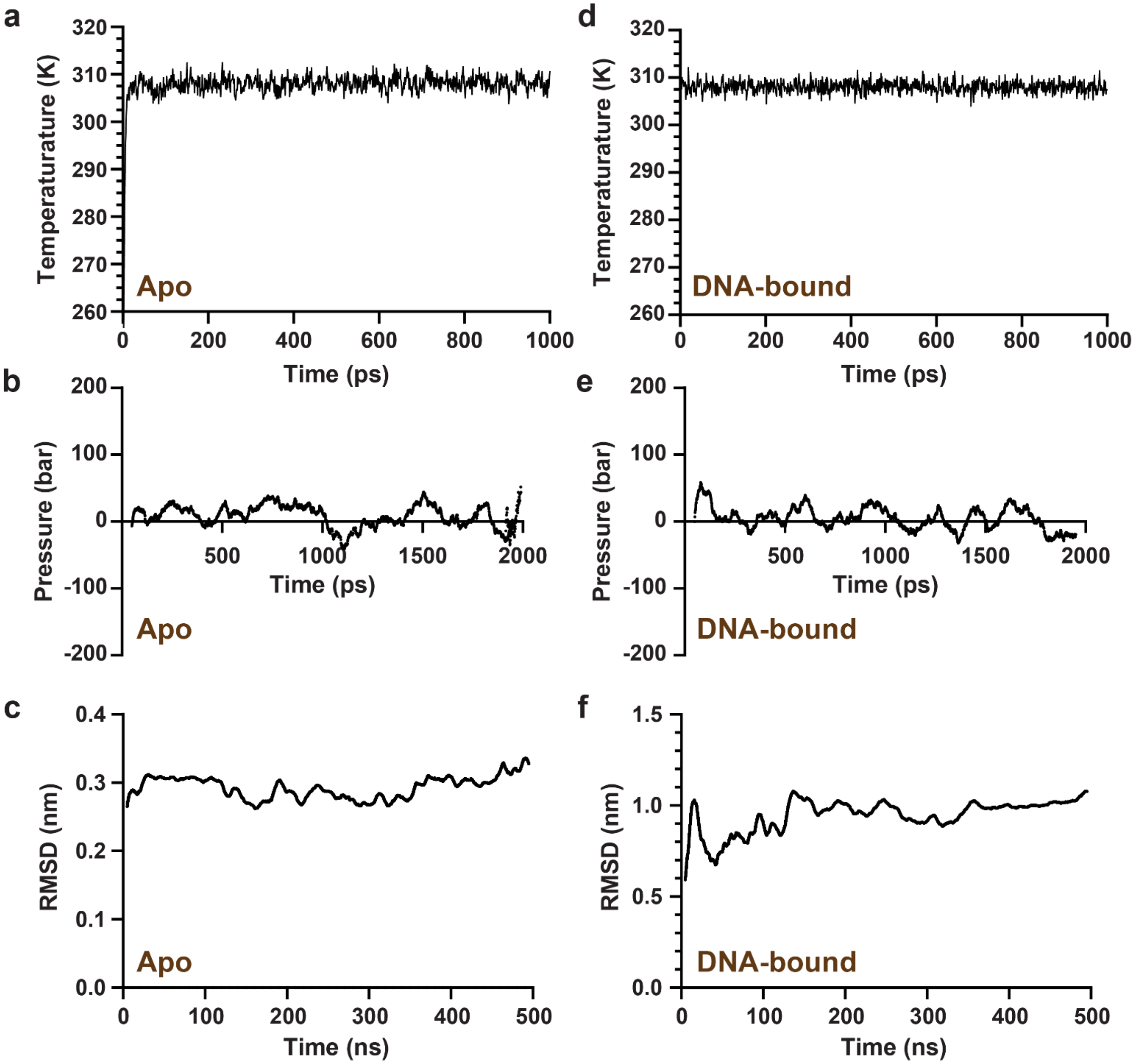


**Extended Data Fig. 4 | Analysis of the MD simulations during different phases**

**a,d,** Equilibration of temperature under the number of particles, volume, and temperature (NVT ensemble) conditions for (**a**) apo and (**d**) DNA-bound CTD.

**b,e,** Equilibration of pressure under the number of particles, pressure, and temperature (NPT ensemble) conditions for (**b**)apo and (**e**) DNA-bound CTD.

**c,f,** RMSD analysis of the MD production phase for (**c**) apo and (**f**) DNA-bound CTD.

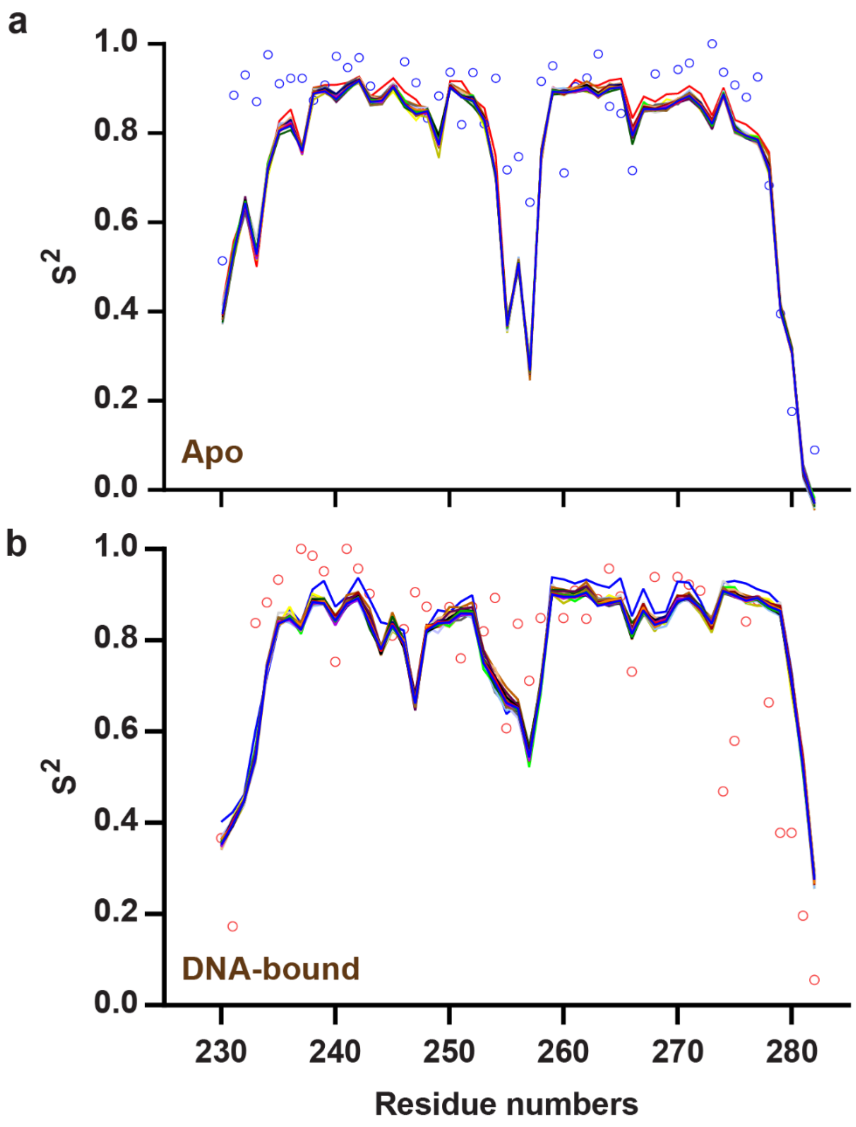


**Extended Data Fig. 5 | Generalized order parameter ($S^2$) comparison**

**a,** Experimental (model-free values) versus computer-simulated (from 500 ns MD trajectories with averaging time windows from 100 ps to 5 ns) $S^2$ values for apo CTD. The open circles represent experimental model-free $S^2$ values, and the solid lines represent molecular dynamics simulation results.

**b,** Those for DNA-bound CTD.

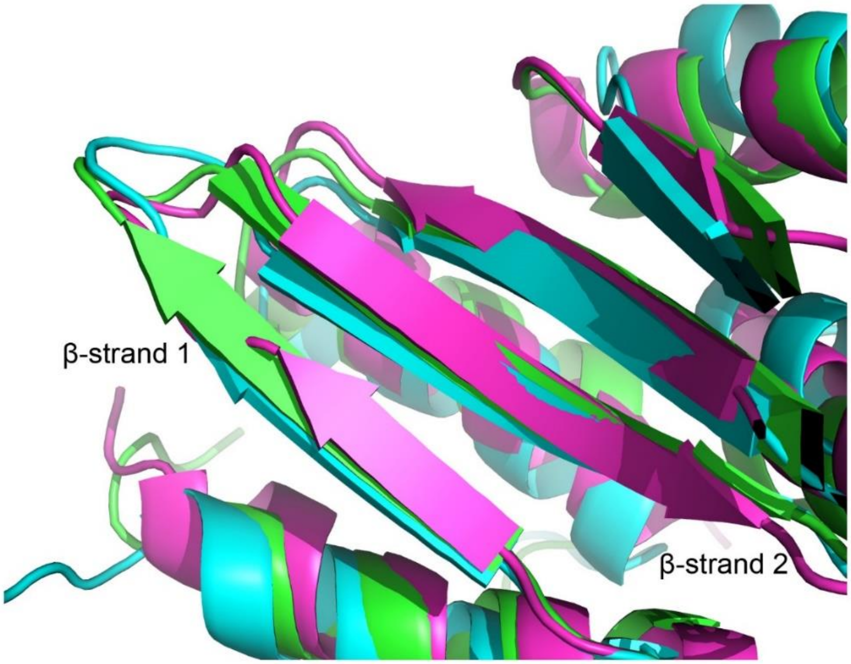


**Extended Data Fig. 6 | An overlay of the ParB C-terminal domain (CTD) structures.**
Average simulated structures of apo (green) and DNA-bound CTD (cyan), along with apo NMR structure (PDF entry: 5NOC) (purple), are overlaid to highlight differences in β-strands. The NMR structure of DNA-bound CTD is not available.

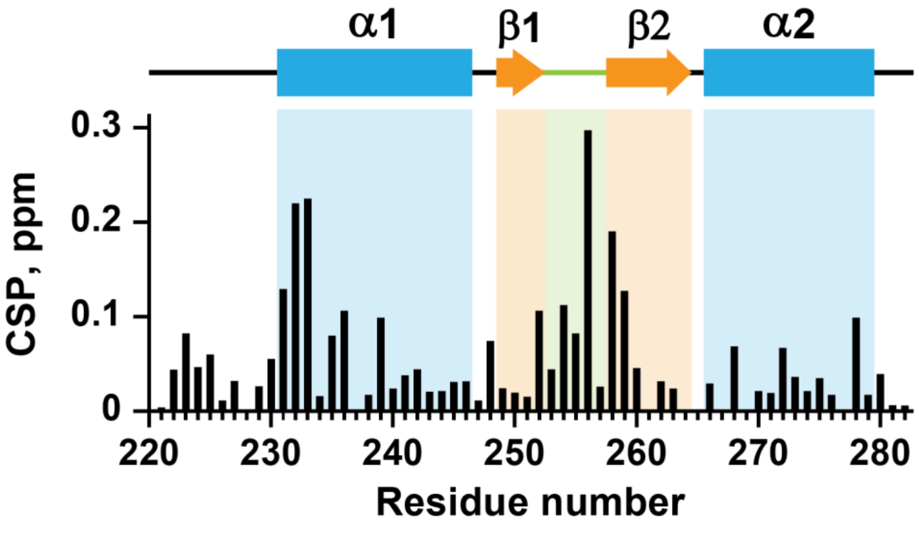


**Extended Data Fig. 7 | Chemical shift perturbation (CSP) of CTD upon addition DNA, at pH 6.1, 0.4 mM.** The secondary structure of the apo CTD determined from the NMR structure (PDB entree 5NOC) is displayed at the top: α-helices (blue bars), β-strands (orange arrows), loop regions (black lines), and linker region (green).
